# The Mutational Landscape of the SCAN-B Real-World Primary Breast Cancer Transcriptome

**DOI:** 10.1101/2020.01.30.926733

**Authors:** Christian Brueffer, Sergii Gladchuk, Christof Winter, Johan Vallon-Christersson, Cecilia Hegardt, Jari Häkkinen, Anthony M. George, Yilun Chen, Anna Ehinger, Christer Larsson, Niklas Loman, Martin Malmberg, Lisa Rydén, Åke Borg, Lao H. Saal

**Affiliations:** Division of Oncology, Department of Clinical Sciences, Lund University, Lund, Sweden; Lund University Cancer Center, Medicon Village, Lund, Sweden; CREATE Health Strategic Center for Translational Cancer Research, Lund University, Lund, Sweden; Department of Pathology, Skåne University Hospital, Lund, Sweden; Division of Molecular Pathology, Department of Laboratory Medicine, Lund University, Lund, Sweden; Department of Oncology, Skåne University Hospital, Lund, Sweden; Department of Surgery, Skåne University Hospital, Lund, Sweden

## Abstract

Breast cancer is a disease of genomic alterations, of which the complete panorama of somatic mutations and how these relate to molecular subtypes and therapy response is incompletely understood. Within the Sweden Cancerome Analysis Network–Breast project (SCAN-B; ClinicalTrials.gov NCT02306096), an ongoing study elucidating the tumor transcriptomic profiles for thousands of breast cancers prospectively, we developed an optimized pipeline for detection of single nucleotide variants and small insertions and deletions from RNA sequencing (RNA-seq) data, and profiled a large real-world population-based cohort of 3,217 breast tumors. We use it to describe the mutational landscape of primary breast cancer viewed through the transcriptome of a large population-based cohort of patients, and relate it to patient overall survival. We demonstrate that RNA-seq can be used to call mutations in important breast cancer genes such as *PIK3CA*, *TP53*, and *ERBB2*, as well as the status of key molecular pathways and tumor mutational burden, and identify potentially druggable genes in 86.8% percent of tumors. To make this rich and growing mutational portraiture of breast cancer available for the wider research community, we developed an open source web-based application, the SCAN-B MutationExplorer, accessible at http://oncogenomics.bmc.lu.se/MutationExplorer. These results add another dimension to the use of RNA-seq as a potential clinical tool, where both gene expression-based and gene mutation-based biomarkers can be interrogated simultaneously and in real-time within one week of tumor sampling.

## Introduction

Mutations in the cancer genome, including single nucleotide variants (SNVs) and small insertions and deletions (indels), can shed light on cancer biology, tumor evolution and susceptibility or resistance to therapeutic agents [1, 2, 3]. Mutations can now even be used to track circulating tumor DNA in the blood of patients [4, 5]. In recent years, the characterization of the mutational landscape of breast cancer has been performed primarily on the DNA level [1, 6, 7]. Adoption of massively parallel RNA sequencing (RNA-seq) as a clinical tool has been slower, despite a number of complementary advantages over DNA-seq. In addition to gene and isoform expression profiling and detection of *de novo* transcripts such as fusion genes, RNA-seq can approximate classical DNA-seq capabilities in the detection of SNVs, indels, as well as structural variants [8] and coarse copy number [9]. This makes RNA-seq an excellent tool for biomarker development [10] and potential clinical deployment [11, 12].

For these reasons, among others, in 2010 the Sweden Cancerome Analysis Network–Breast (SCAN-B) initiative (ClinicalTrials.gov ID NCT02306096) selected RNA-seq as the primary tool [13, 14]. SCAN-B is a prospective real-world and population-based multicenter study with the aim of developing, validating, and clinically-implementing novel biomarkers. To this end, SCAN-B collects tumor tissue and blood samples from enrolled patients with a diagnosis of primary breast cancer (BC). To date over 14,000 patients have been enrolled, and messenger RNA (mRNA) sequencing is performed on patient tumors within one week of surgery. All patients are treated uniformly according to the Swedish national standard of care regimen.

Expression profiling is an excellent tool to develop gene signatures for established and novel biomarkers [10, 15, 16], and many such signatures can be applied to a single RNA-seq dataset. However, for the detection of SNVs and indels from RNA-seq data there are a number of challenges. Unlike DNA-seq, where whole-genome or targeted sequencing reads are distributed approximately uniformly and in proportion to DNA copy number, the abundance of reads in RNA-seq is proportional to the expression of each gene or locus. Consequently, only variants in expressed transcripts of sufficient level can be detected. In cancer, this means that variants in oncogenes can likely be detected, whereas those in tumor suppressor genes, e.g. *TP53*, *BRCA1*, or *BRCA2*, are more likely to be missed. For example, mutations inducing premature stop codons can lead to nonsense-mediated decay, causing loss of expression and subsequently false-negative calls. The transcriptome is also more complex and challenging than the genome. RNA structures, such as alternative splicing, add computational challenges to alignment, and RNA editing can contribute to false-positive variant calls. Finally, there is the lack of benchmark datasets for RNA-seq, as are available for DNA from the Genome in a Bottle consortium and others [17, 18].

The aim of this study was to optimize RNA-seq somatic mutation calling through comparison to matched targeted DNA-seq, discern the mutational landscape of the early breast cancer transcriptome across a large cohort of 3,217 treatment-naïve SCAN-B cases with sufficient follow-up time, and to make the resulting vast dataset available for exploration by the wider research community. To demonstrate the power of the methodology and dataset, we assessed the impact of mutations in important breast cancer driver genes and pathways, as well as tumor mutational burden (TMB) on patient overall survival (OS).

## Materials and Methods

### Patients

The study was approved by the Regional Ethics Review Board of Lund at Lund University (diary numbers 2007/155, 2009/658, 2009/659, 2010/383, 2012/58, 2013/459). We analyzed data from two previously described cohorts. For 273 patients, including two patients with bilateral disease (thus 275 tumors), enrolled in the All Breast Cancer in Malmö (ABiM) study from 2007-2009, matched snap-frozen primary breast tumor tissue and blood samples were collected as previously described [19]. A cohort of 3,273 SCAN-B primary breast tumors described previously [10] was reduced to 3,217 samples following additional quality controls. Tissue collection, preservation in RNAlater, sequencing, expression estimation, and molecular subtyping using the PAM50 gene list were performed as previously reported [10, 13]. Clinical records were retrieved from the Swedish National Cancer Registry (NKBC). Estrogen receptor (ER) and Progesterone receptor (PgR) status was categorized using an immunohistochemical staining cutoff of 1%. Patients in the SCAN-B cohort had median 74.5 months follow-up, and patient demographics for both cohorts are detailed in Table 1.

### Library Preparation and Sequencing

For the 275 sample ABiM cohort, tumor and normal DNA was sequenced using a custom targeted capture panel of 1,697 genes and 1,047 miRNAs as described [19]. For the same tumors RNA-seq was performed as described [10] (a subset of the 405 sample cohort therein). In short, strand-specific dUTP libraries were prepared and sequenced on an Illumina HiSeq 2000 sequencer to an average of 50 million 101bp reads per sample [13, 20].

For the 3,217 sample SCAN-B cohort, RNA-seq data was generated as previously described [10]. In short, strand-specific dUTP mRNA-seq libraries were prepared [13, 20], and an average 38 million 75bp reads were sequenced on an Illumina HiSeq 2000 or NextSeq 500 instrument (Supplementary Table S1).

### Sequence Data Processing

For tumor and normal DNA, reads were aligned to the hg19 reference genome using Novoalign 2.07.18 (Novocraft Technologies, Malaysia). Using a modified version of the variant workflow of the bcbio-nextgen NGS framework (https://github.com/bcbio/bcbio-nextgen, modified version https://github.com/cbrueffer/bcbio-nextgen/tree/v1.0.2-scanb-calling) utilizing Bioconda for software management [21], duplicate reads were marked using biobambam v2.0.62 [22] and variants were called from paired tumor/normal samples using VarDict-Java 1.5.0 [23] (with default options except `-f 0.02 -N ${SAMPLE} -b ${BAM_FILE} -c 1 -S 2 -E 3 -g 4 -Q 10 -r 2 -q 20`), which internally performs local realignment around indels. Variant coordinates were converted to the GRCh38 reference genome using CrossMap 2.5 [24].

Raw RNA-seq reads were trimmed and filtered as described previously [10], and then processed using the modified bcbio-nextgen 1.0.2 variant workflow. Reads were aligned to a version of the GRCh38.p8 reference genome that included alternative sequences and decoys, and was patched with dbSNP Build 147 common SNPs, and the GENCODE 25 transcriptome model using HISAT2 2.0.5 [25] (with default options except `--rna-strandness RF --rg-id ${ID_NAME} --rg PL:illumina --rg PU:${UNIT} --rg SM:${SAMPLE}`). BAM index files were generated using Sambamba 0.6.6 [26] and duplicate reads were marked using SAMBLASTER 0.1.24 [27]. Variants were called using VarDict-Java 1.5.0 with default options except `-f 0.02 -N ${SAMPLE} -b ${BAM_FILE} -c 1 -S 2 -E 3 -g 4 -Q 10 -r 2 -q 20` callable_bed, where callable_bed was a sample-specific BED file containing all regions of depth *≥* 4.

All variants were annotated using a Snakemake [28] workflow around vcfanno 0.3.1 [29] and the data sources dbSNP v151 [30], Genome Aggregation Database (gnomAD) [31], Catalogue of Somatic Mutations in Cancer (COSMIC) v87 [32, 33], CIViC [34], MyCancerGenome (release March 2016, http://www.mycancergenome.org), SweGen version 20171025 [35], the Danish Genome Project population reference [36], RNA editing databases [37, 38, 39, 40], UCSC low complexity regions, IntOGen breast cancer driver gene status [41] (accessed 2018-08-02), and the drug gene interaction database (DGIdb) v3.0.2 [42]. We used SnpEff v4.3.1t (with default parameters except `hg38 -t -canon`) [43] to predict functional variant impact on canonical transcripts as defined by SnpEff.

To filter out recurrent artifacts introduced during library preparation or sequencing, we constructed a panel of “normal” tissues consisting of all variants enumerated from RNA-seq analysis of adjacent non-tumoral breast tissues sampled from ten SCAN-B patients.

Gene expression data in fragments per kilobase of transcript per million mapped reads (FPKM) for the ABiM cohort was generated as previously reported [10] and is available from the NCBI Gene Expression Omnibus (accession number GSE96058).

### Variant Filtering

The strategy we applied for developing DNA-seq-informed filters is outlined in Figure 1. Due to the lenient settings used for sensitive initial variant calling, we developed and applied rigid filters to reduce false positive calls resulting from either sequencing or PCR artifacts, RNA editing, or germline variants. To this end, variants called from 275 matched tumor/normal targeted capture DNA datasets were filtered, among other parameters, for low complexity regions, SNP status (dbSNP “common”, SweGen and COSMIC SNPs, high gnomAD allele frequency), allele frequency ≥ 0.05, depth ≥ 8, homopolymer environments, and RNA editing sites. Using the resulting DNA variants as reference we developed filters for the 275 sample RNA-seq variants by permuting values of the sequencing, variant calling, and annotation variables, and for each permutation calculating the concordance to the DNA mutations. Following these “negative” filters we applied a range of “positive” filters to rescue filtered variants, e.g. to retain a variant if it is present in the curated MyCancerGenome database of clinically important mutations. Finally, we selected the combination of “negative” and “positive” filter settings with the best balance of sensitivity and specificity. Using SnpSift [44] we applied the filters to RNA-seq mutation calls from the 3,217 patient cohort. A complete list of final filter variables and values for both the tumor/normal DNA variant calls, as well as the RNA-seq variant calls can be found in Supplementary Table S2.

**Figure 1:**
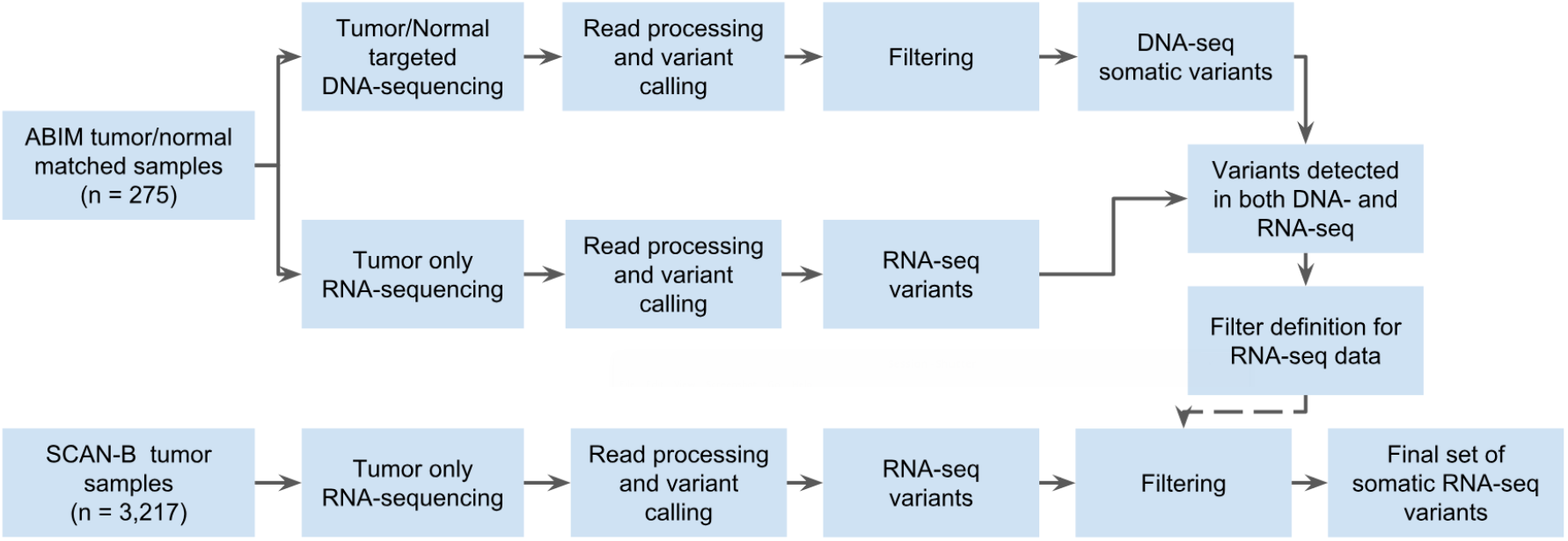
Study design flow diagram for DNA-seq-informed optimization of RNA-seq variant calling.

### Data Analysis

All analyses were performed using R 3.5.1. Waterfall and lollipop plots were made using the *GenVisR* 1.14.2 [45] and *RTrackLayer* 1.42.1 packages. Substitution signatures were analyzed using the *MutationalPatterns* 1.8.0 package [46]. Survival analysis was conducted using OS as endpoint. Overall survival was analyzed using the Kaplan-Meier (KM) method and two-sided log-rank tests implemented in the *survival* 2.44-1.1 package. Associations were tested using Fisher’s exact test. P-values *<* 0.05 were considered significant. The web application SCAN-B MutationExplorer was written in R using the Shiny, GenVisR and SurvMiner packages.

## Results

### Variant Filter Performance

Mutation calling in the 275 sample cohort resulted in 3,478 somatic post-filter mutations from the matched tumor/normal targeted capture DNA, and 1,459 variants from tumor RNA-seq in the DNA capture regions (Table 2 and Supplementary Figure S1). Comparing these DNA and RNA variants resulted in 1,132 mutations that were present both in DNA and RNA in the capture regions and whose frequencies were generally in line with previous studies such as The Cancer Genome Atlas (TCGA) [1] (Supplementary Figure S2). Of the 1,459 RNA-seq variants, 884 (60.6%) were identified as somatic in DNA, 248 (17.0%) as germline in DNA, and 327 (22.4%) as unique to RNA. These RNA-unique variants are a mix of somatic mutations missed in DNA-seq, e.g. due to regional higher sequencing coverage in RNA-seq or tumor heterogeneity, unfiltered RNA-editing sites, or artifacts caused by PCR, sequencing, or alignment and variant calling.

### Landscape of Somatic Mutations in the Breast Cancer Transcriptome

We applied the filters derived from the 275 sample set to the entire RNA-seq SCAN-B 3,217 sample set, resulting in 144,593 total variants comprised of 141,095 SNVs, 1,112 insertions, and 2,386 deletions (Table 2). The number of per-sample mutations in the SCAN-B set was lower compared to the ABiM set, likely due to the ABiM set being sequenced to a higher depth (Supplementary Table S1). The SNVs comprised 50,270 missense, 2,311 nonsense, 1,042 splicing, 68,819 affecting 3’/5’ untranslated regions (UTRs), 17,057 synonymous mutations, as well as 1,596 mutations predicted otherwise. The majority of indels were predicted to cause frame-shifts, or affect 3’/5’ UTRs (Supplementary Table S3). After removing synonymous mutations the number of mutations was reduced to 127,536 variants in the SCAN-B set, i.e. an average of 40 mutations per tumor.

We analyzed the contribution of the six nucleotide substitution types (`C>A`, `C>G`, `C>T`, `T>A`, `T>C`, and `T>G`) to SNVs in the ABiM and SCAN-B sets (Figure 3B). Compared to DNA, RNA-seq based variant calls showed a relative under-representation of `C>T` substitutions, and an over-representation of `T>C` substitutions.

**Figure 3:**
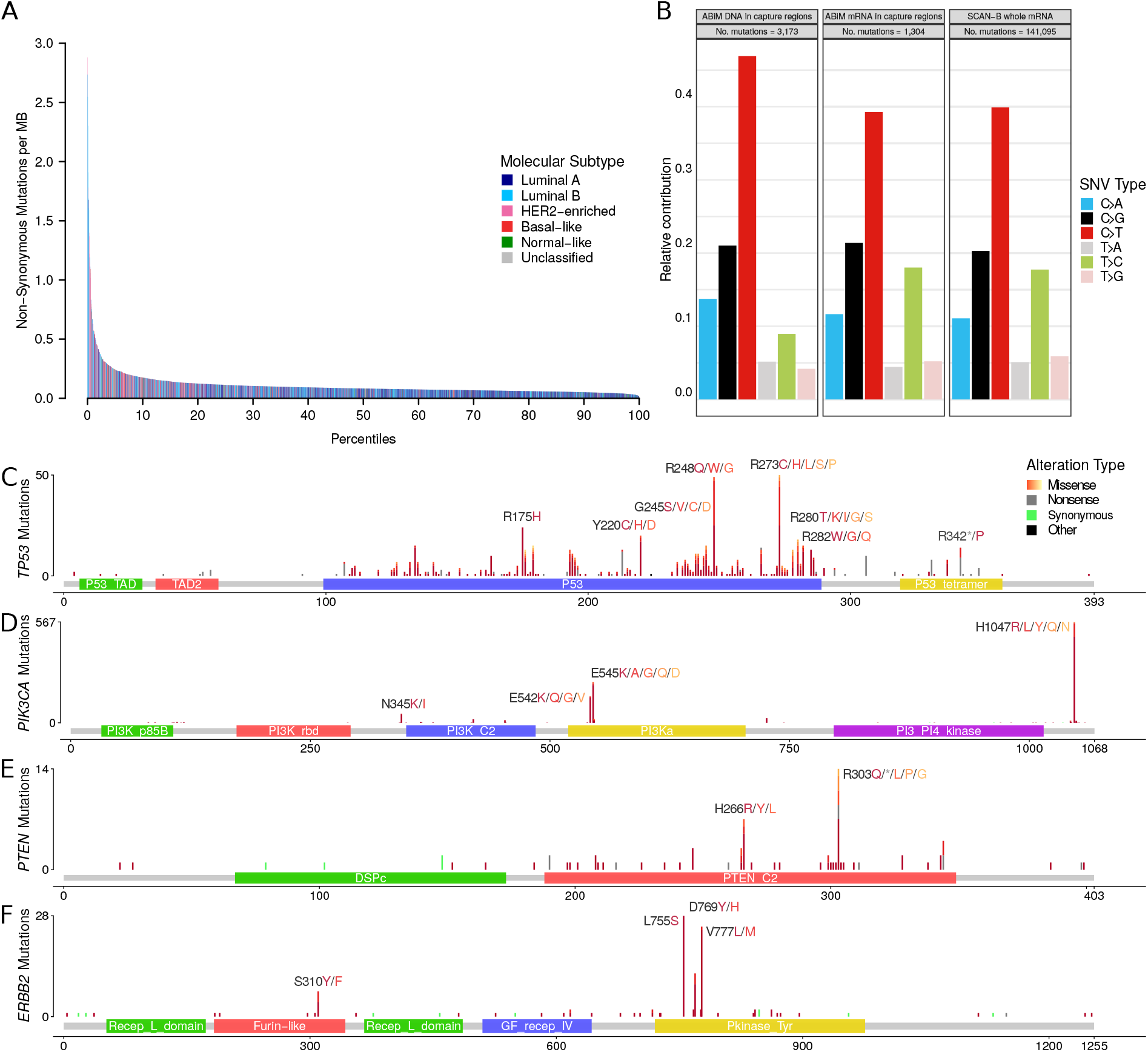
**A**) Number of non-synonymous mutations per sample. Bars are colors by PAM50 subtypes Luminal A (dark blue), Luminal B (light blue), HER2-enriched (pink), basal-like (red), Normal-like (green) and Unclassified (gray). **B**) Contribution of base change types to the overall SNV composition in the ABiM cohort for captured DNA regions and mRNA in the captured DNA regions, as well as SCAN-B whole mRNA. **C-F**) Lollipop plots showing the abundance, location, and type of SNVs in the genes **C**) *TP53*, **D**) *PIK3CA*, **E**) *PTEN*, and **F**) *ERBB2*. Protein change labels are shown for the most mutated amino acid positions, with residues ordered left to right by mutation frequency.

In accordance with published studies of primary BC, the most frequently mutated genes were the known BC drivers *PIK3CA* (34% of samples), *TP53* (23%), *MAP3K1* (7%), *CDH1* (7%), *GATA3* (7%), and *AKT1* (5%) (Figure 2). As reported before [6], disruptive alterations in *CDH1* were a hallmark of lobular carcinomas (135/386 [35.0%] of samples), while alterations in *TP53*, *MAP3K1*, and *GATA3* were more common in the ductal type. 86.8% of SCAN-B samples had at least one mutation in a gene targeted by an approved or experimental drug, based on the Database of Gene-drug Interactions (DGI).

**Figure 2:**
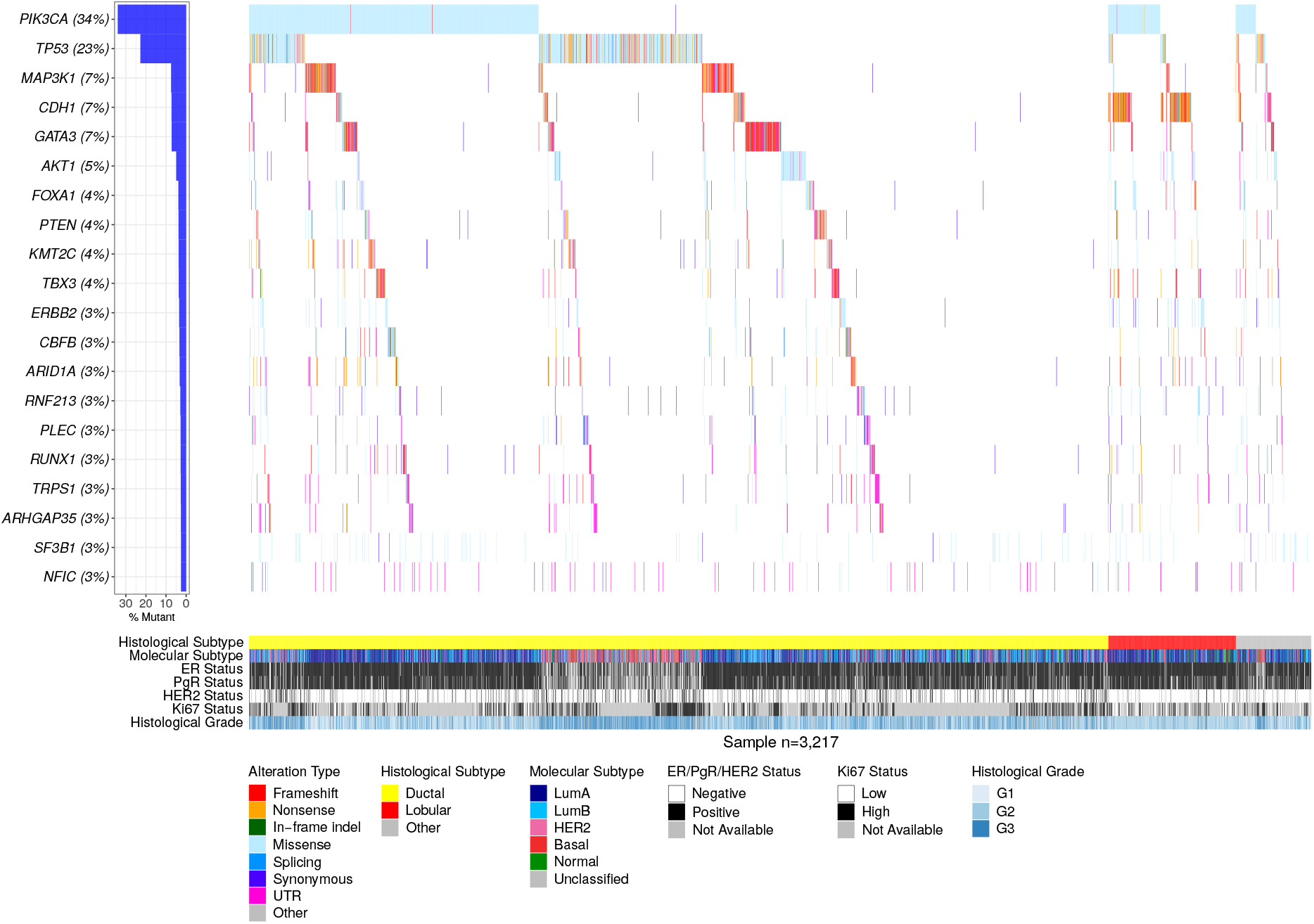
Waterfall plot of the 20 most frequent post-filter RNA variants across 3,217 SCAN-B samples. Genes are ranked by variant frequency in RNA-seq. Samples are sorted by histological subtype and alteration occurrence.

### Somatic Mutations in Important BC Genes

We looked closer into known driver BC genes and found our RNA-seq based mutation calls to recapitulate known mutation rates and hotspots, summarized in Table 3, Supplementary Table S3, and Figure 3C-F. Associations of mutated genes and clinical and molecular biomarkers are summarized in Table 4, and we highlight several examples below.

*PIK3CA* was the most frequently mutated gene, with 1,163 non-synonymous mutations in 1,095 patient samples (34% of patients). As expected and in line with previous studies [1, 47] the majority of alterations were the known hotspot mutations H1047R/L, E545K, and E542K (Table 3, Figure 3D). All of these hotspot mutations and the vast majority of other *PIK3CA* alterations were missense mutations. Mutations were associated with lobular, ER+, PgR+, HER2-, and Luminal A (LumA) BC (Table 4).

*TP53* is frequently disrupted by somatic SNVs, however a few hotspot mutations exist [48]. The mutation frequency in BC estimated to be 35.4%-37% [1, 47], which we could confirm in our DNA-seq ABiM filter definition cohort (37%). Likely due to nonsense-mediated decay, loss of heterozygosity, and/or decreased mRNA transcription, in the 3,217 cases the frequency of *TP53* mutations was lower at 23% (782 mutations in 733 samples). Nevertheless, as anticipated *TP53* mutations were associated with ductal, ER-, PgR-, HER2+, triple-negative BC (TNBC), and the basal-like and HER2-enriched PAM50 subtypes (Table 4). The most often mutated amino acids we observed were R273, R248, R175 (50, 49, and 24 mutations respectively, total 123/782 [15.7%]), followed by positions Y220 (21/782 [2.7%]), R280 (19/782 [2.4%]) and R342 (17/782 [2.2%]), in line with the IARC *TP53* database (release R19) [49] (Table 3, Figure 3C).

*PTEN* is a crucial tumor suppressor gene and regulator of PI3K activity, and PTEN protein expression is associated with poor outcome [50]. In our dataset we found 124 non-synonymous mutations in 116/3,217 (3.6%) samples, and mutations were significantly associated with HER2-disease (Table 4).

We identified 117 non-synonymous *ERBB2* mutations in 103 patients (3.2%), higher than the previously reported incidence rates of 1.6%-2.4% [2, 51, 52], but lower than in metastatic BC where rates as high as ~7% have been reported [53]. Two hotspots, L755S (28/117) and V777L (24/117) that cause constituent HER2 signaling (Figure 3F) [2, 51], accounted for 44.4% of total *ERBB2* mutations. Co-occurrence of *ERBB2* mutation and amplification has been reported before, however mainly in the metastatic setting and after treatment [53]. In our untreated, early BC cohort we observed *ERBB2* mutation and amplification in 12 tumors, and generally mutation and amplification were not mutually exclusive (P=0.88), and *ERBB2* mutations were predominantly in tumors classified as PAM50 HER2-enriched subtype (P=0.001). Moreover, *ERBB2* mutation was significantly associated with PgR-, and lobular BC (Table 4)

Loss of E-cadherin (CDH1) protein expression is a hallmark of the lobular BC phenotype [6]. With 12% of our cohort being of lobular type, we observed 137 of total 233 *CDH1* mutations in lobular BCs (58.8%, P<0.001). The mutations were mostly comprised of nonsense mutations (37.2%) and frameshift indels (35.4%), suggesting they lead to CDH1 expression loss and drive the lobular phenotype. We observed one nonsense mutation hotspot (Q23*, n=18), and this residue was also hit by a rare missense mutation (Q23K, n=1). In addition to lobular BC, *CDH1* mutations were associated with ER+/HER2-status, and the LumA subtype (Table 4).

Other notable mutated genes in our set were *MAP3K1*, *AKT1*, *ESR1*, *GATA3*, *FOXA1*, *SF3B1*, and *CBFB*. *MAP3K1* is a regulator of signaling pathways, and regularly implicated in various cancer types. We observed a high rate of frame-shift indels, and missense mutations mostly clustered in the kinase domain. Co-mutation of *MAP3K1* and *PIK3CA* occurred in 108 tumors (3.4%). *AKT1* is a common oncogene with 156 (4.8%) mutated samples, and featured the fourth most mutated hotspot (E17K, 121 mutations) in the SCAN-B cohort. These mutations are predictive of sensitivity to AKT inhibitors [54]. *ESR1* encodes the ER*α* receptor, perhaps the most important clinical BC biomarker, and 77 tumors harbored 81 *ESR1* variants. Relatedly, *GATA3* and *FOXA1* are frequently mutated transcription factors involved in ER signalling. We identified 246 *GATA3* mutations, including known recurrent frame-shift mutations (P409fs, n=30 and D336fs, n=10) and the M294K/R missense mutation (n=15), as well as 10 splice site variants. In *FOXA1* we detected 146 total mutations, including known recurrent S250F (n=23) and F266L/C (n=12) missense mutations. Most mutations occurred in the forkhead DNA binding domain. In line with their involvement in ER signalling, mutations in *GATA3*, *FOXA1*, *MAP3K1*, and *ESR1* were associated with ER+ and PgR+ disease. Further, *GATA3* and *MAP3K1* were associated with ER2+/HER2- and ductal BC, while *ESR1* and *FOXA1* were more common in lobular BC. With the exception of *GATA3* (Luminal B [LumB]), all of these genes were associated with the LumA subtype (Table 4). In *SF3B1* we identified 81 mutations in 79 tumors, 60 of which were K700E hotspot mutations. Alterations in this gene are associated with ER+ disease [55] and affect alternative splicing patterns [56]. The cohort frequency of 1.9% K700E mutations matches up with previously reported 1.8% in an unselected breast cancer cohort [55]. We could confirm the reported prevalence of *SF3B1* mutations in ER+ tumors (76/79 mutations in ER+ tumors, P=0.021), as well as the association with non-ductal, and non-lobular subtypes (P=0.003). *CBFB* has been shown to be commonly mutated in ER+/HER2-disease, however the significance of these mutations is unknown [57]. We could confirm this finding showing 107 mutations (3.3% cohort frequency), 95 of which were in ER+/HER2-samples (4% of ER+/HER2-samples, P<0.01). We also found them to be associated with the LumA subtype (Table 4).

### Mutations in Molecular Pathways

We were interested whether the mutational data, when considered from the perspective of mutated pathways, could reveal new biological correlates. To test this, we mapped mutation status to important BC pathways as defined in the Reactome database [58, 59]. We called a pathway mutated when at least one of the member genes had a non-synonymous mutation, and clustered samples by pathway mutation status using Ward linkage. Notable clusters that emerged were co-mutated hedgehog signaling, p53-independent DNA repair, and hypoxia response pathways, as well as a cluster of NOTCH1/2/3 signaling mutated tumors, both in mostly basal-like and HER2-enriched tumors, and PI3K/AKT, MET, RET, EGFR, ERBB2, and ERBB4 signaling pathways co-mutated in a subset of tumors of Luminal A and B subtype (Figure 4, see Supplementary Table S4 for Reactome pathway IDs).

**Figure 4:**
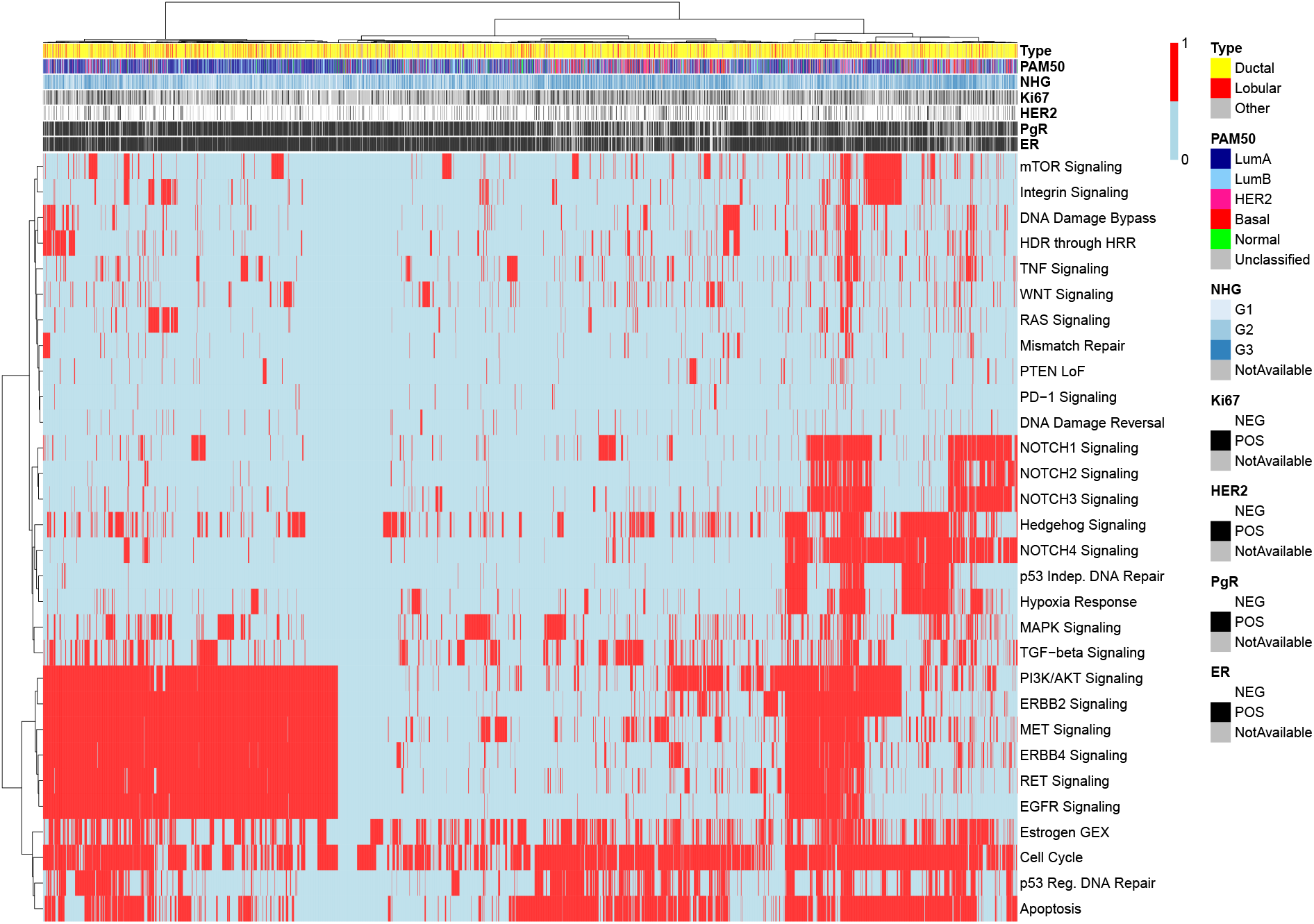
Binary heatmap of mutation status of important BC pathways in 3,217 samples. Samples with wild-type (wt) pathway status (all member genes wt) are colored blue, those with mutated pathways (at least one member gene mutated) are colored red. Reactome IDs for the pathways can be found in Supplementary Table S4.

### Tumor Mutational Burden

Tumor mutational burden is increasingly of interest due to its association to neoantigen burden and response to immunotherapies. We used the median number of non-synonymous mutations per transcriptome megabase (rnaMB), 0.082 mutations/rnaMB, to stratify all SCAN-B samples into TMB-high and TMB-low groups. Samples with HER2-enriched and basal-like PAM50 subtypes were enriched in the top 10% of samples with the highest TMB compared to the lowest 90% (P<0.001, Figure 3A), supporting previous results and indicating that immunotherapy may have higher activity in these two PAM50 subtypes [1].

### Mutational Landscape and Patient Outcomes

Next, we were interested in the association between mutations in important BC genes and patient outcome under various treatments. Below we show the results for *TP53*, *PIK3CA*, *ERBB2*, and *PTEN* with OS of SCAN-B patients in clinical biomarker and treatment groups (Figure 5, Supplementary Figure S4), as well as selected pathways (Figure 6, Supplementary Figure S5). The web tool SCAN-B MutationExplorer may be used to query any gene(s) of interest.

**Figure 5:**
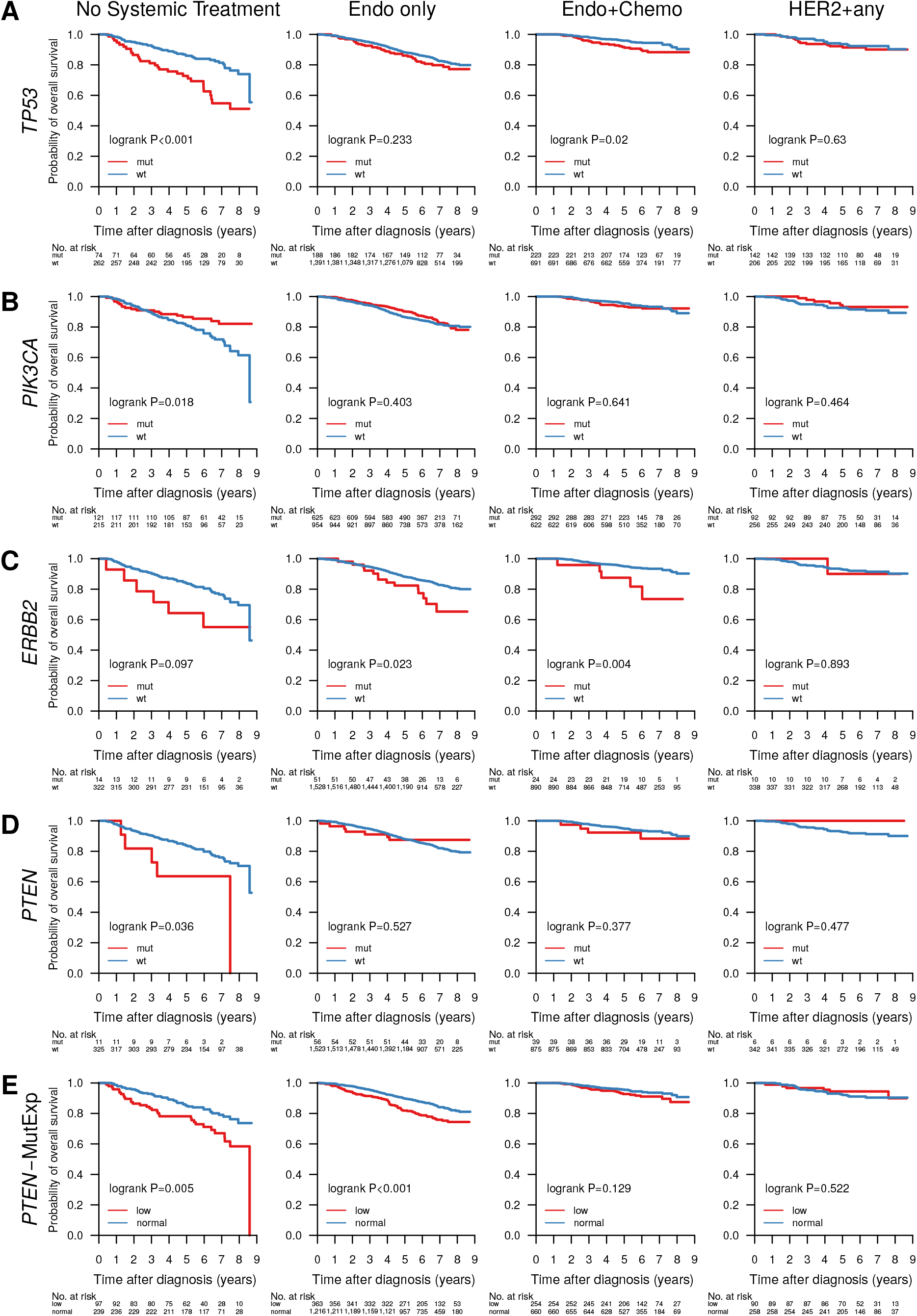
Overall survival of patients with tumors containing mutations in the genes **A**) *TP53*, **B**) *PIK3CA*, **C**) *ERBB2*, **D**) *PTEN*, and **E**) mutated *PTEN* or *PTEN* expression in the lower quartile across the cohort, stratified by groups receiving no systemic treatment, endocrine therapy only (Endo only), endocrine and chemo therapy (Endo+Chemo), as well as HER2 treatment with any other treatment or none (HER2+any).

**Figure 6:**
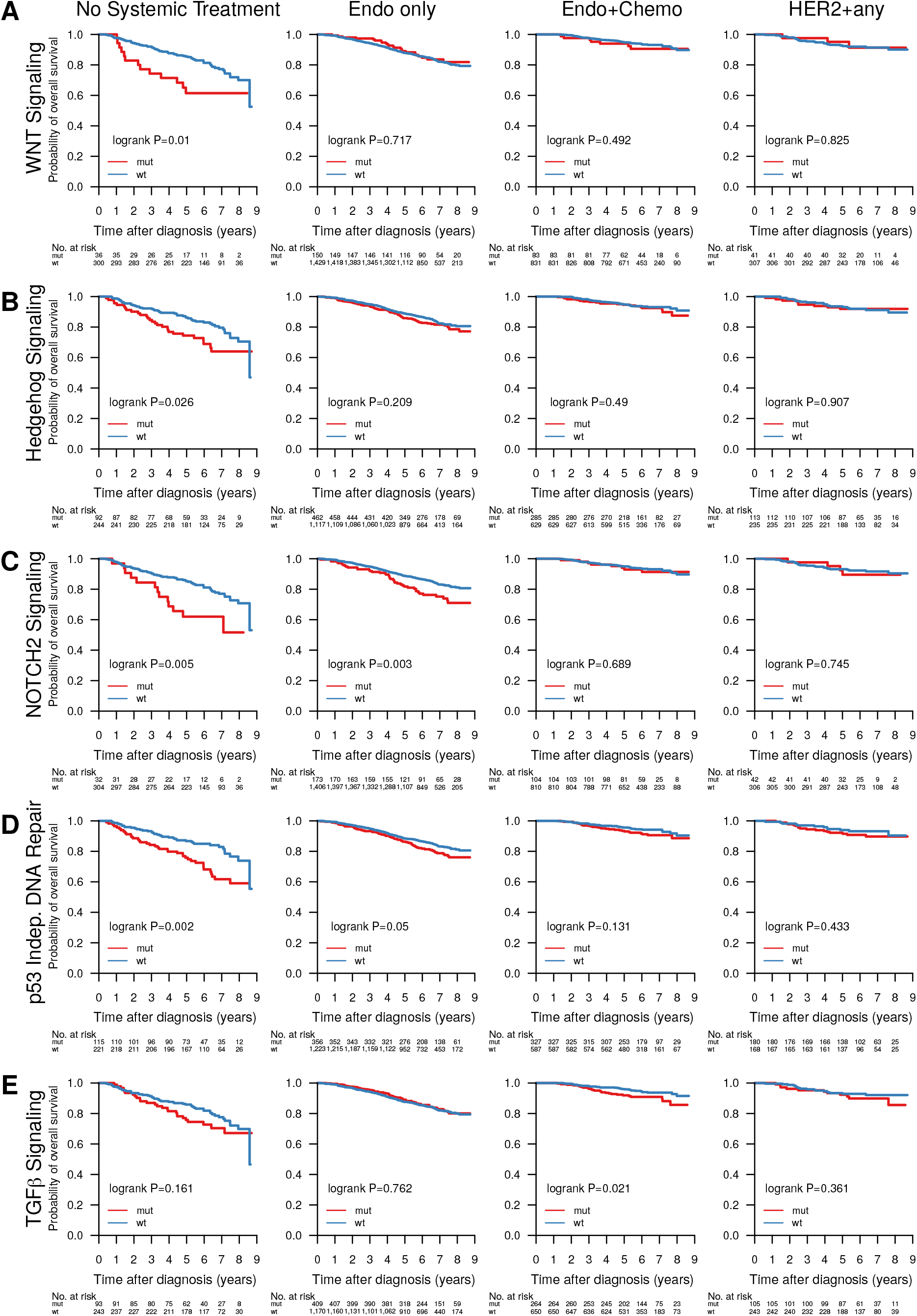
Overall survival of patients with tumors containing mutations in pathways **A**) WNT signaling, **B**) Hedgehog signaling, **C**) NOTCH2 signaling, **D**) p53 independent DNA damage repair, **E**) TGF*β* signaling, stratified by groups receiving no systemic treatment, endocrine therapy only (Endo only), endocrine and chemo therapy (Endo+Chemo), as well as HER2 treatment with any other treatment or none (HER2+any). See Supplementary Table S4 for Reactome pathway IDs.

In line with expectations, *TP53* mutation predicted poor survival in untreated patients (P<0.001), patients treated with endocrine and chemo therapy (P=0.02), as well as the HR+/HER2-biomarker subgroup (P=0.019).

In early-stage breast cancer *PIK3CA* mutations have been shown to be associated with benefits in disease-free survival [60]. In our hands these mutations signaled worse OS in TNBC patients (P=0.049), and those who did not receive systemic treatment (P=0.018).

*ERBB2* mutations were indicators of poor prognosis in endocrine therapy only (P=0.023) and endocrine and chemo therapy treated (P=0.004) patients, as well as in the ER+/HER2-subgroup (P=0.013).

*PTEN* mutations alone were associated with poor survival in the patient group not receiving systemic treatment (P=0.036), but not in any of the other tested clinical biomarker or treatment groups (Figure 5). While loss of PTEN protein expression or non-functional PTEN protein can be caused by SNVs and indels, it can also be caused by other mechanisms such as large structural variants [61] and promoter methylation [62] that have not been investigated in this study. To account for this we defined a new subgroup *PTEN*-MutExp, where a status of “low” identifies cases with either *PTEN* mutation or gene expression in the lower quartile within the cohort, and “normal” otherwise. The *PTEN*-MutExp low group, incorporating gene expression, showed improved stratification in the no systemic treatment group (P=0.005), and significantly lower OS in patients receiving only endocrine treatment (P<0.001), as well as ER+/HER2-patients (P=0.001). Most of the prognostic value is provided by the gene expression, however mutation data improved stratification (Supplementary Figure S3).

Abstracting from mutations in individual genes we investigated the effect of mutated pathways on OS in patient subgroups stratified by treatment (Figure 6) and clinical subgroup (Supplementary Figure S5). Mutated WNT (Figure 6A, P=0.01), Hedgehog (Figure 6B, P=0.026) and NOTCH2 (Figure 6C, P=0.005) signaling, as well as the p53-independent DNA damage repair (Figure 6D, P=0.002) pathway were associated with worse survival in patients not receiving systemic treatment. Additionally, NOTCH2 signaling (Figure 6C, P=0.003) was associated with worse OS in patients receiving only endocrine treatment, and TGF*β* signaling (Figure 6E, P=0.021) with worse OS in patients treated with endocrine and chemo therapy. Further, WNT signaling was associated with worse OS in HR+/HER2+ (P=0.034) and TNBC patients (P=0.006) (Supplementary Figure S5).

Given its importance as an emerging biomarker for response to immune checkpoint therapy [63], we investigated whether TMB could also provide response information with respect to conventional treatment regimens (Figure 7). When stratified into TMB-high and TMB-low by the SCAN-B cohort median TMB per MB of expressed transcriptome, low TMB was favourable to OS independent of treatment across the cohort (P<0.001), as well as in patients not systemically treated (P<0.001), treated with endocrine therapy only (P<0.001), endocrine ± chemo or HER2 therapy (P=0.003), and chemotherapy ± endocrine or HER2 therapy (P=0.011). High TMB is typically associated with improved survival in TNBC, possibly due to increased neoantigen load enabling a stronger immune response. However, we observed no such effect in TNBC patients within the SCAN-B cohort (P=0.338, Supplementary Figure S6). Mutational load was also a significant survival stratifier across the NHG grading scheme (G1, P=0.022, G2, P=0.008, G3, P=0.005), and within the ER+ (P=0.001), PgR+ (P=0.031), HER2-(P<0.001), and Ki67-high (P=0.006) patient subgroups (Supplementary Figure S6). Interestingly, LumB patients with high TMB showed worse survival (P=0.006), whereas TMB was not a significant stratifier for any other molecular subtype (Supplementary Figure S6).

**Figure 7:**
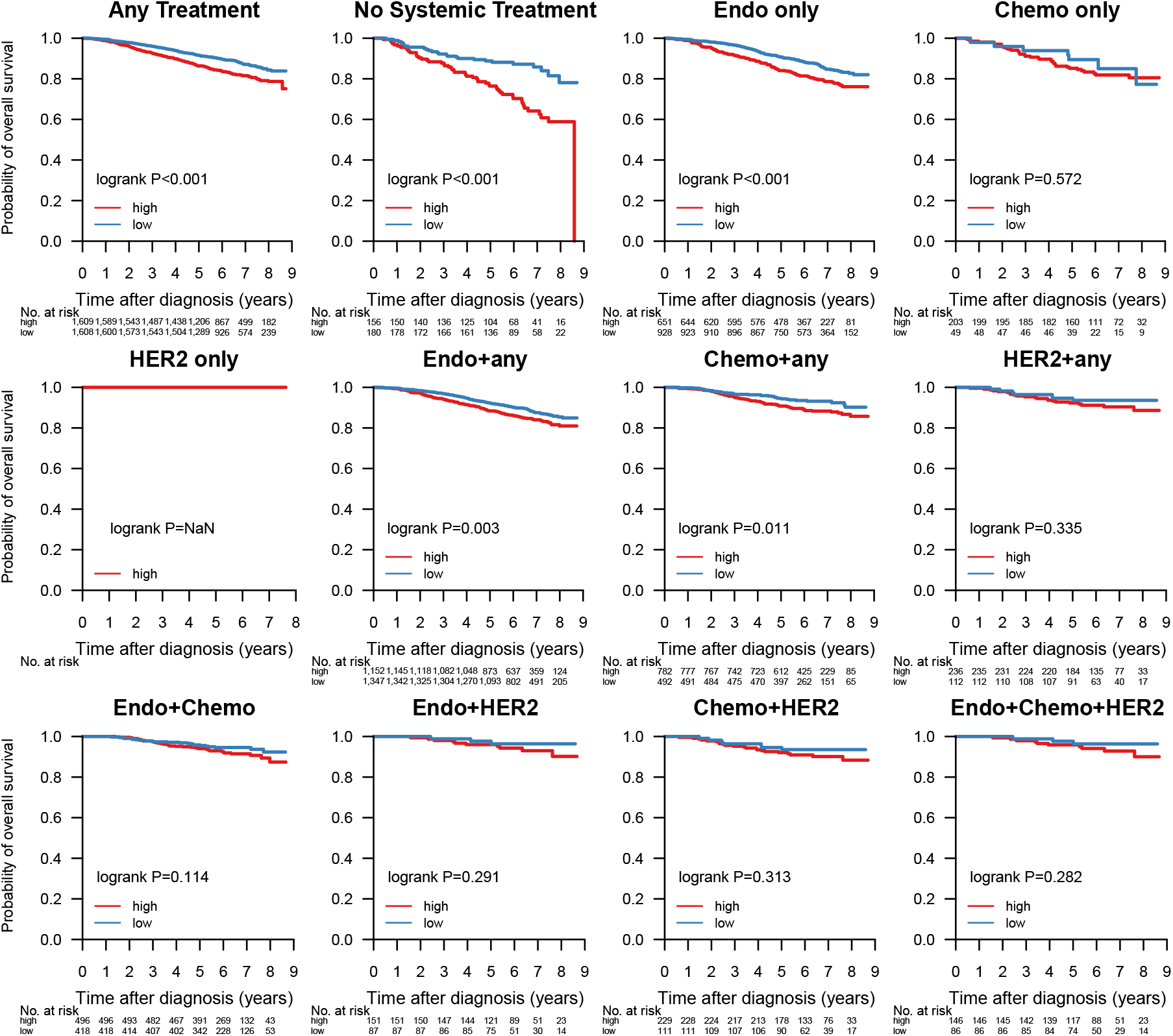
Overall survival stratified by tumor mutational burden (TMB) across different treatment groups. Samples were classified as TMB-high if the amount of non-synonymous mutations per expressed MB (rnaMB) was ≥ the median number of non-synonymous mutations per rnaMB across the whole SCAN-B cohort (0.082 mutations) and TMB-low otherwise.

### SCAN-B MutationExplorer

To enable public exploration and re-use of our rich mutational dataset we developed the web-based application SCAN-B MutationExplorer (available at http://oncogenomics.bmc.lu.se/MutationExplorer) (Figure 8). With this interactive application a user can filter the 3,217 SCAN-B samples based on biomarker combinations (ER, PgR, HER2, Ki67, NHG, histological type, and PAM50 subtype), and mutations based on mutation type (e.g. nonsense or missense) and COSMIC occurrence. From the filtered data the user can create mutational landscape waterfall plots, conduct survival analysis using KM analysis and log-rank tests based on mutations in single genes, pathways as defined in the Reactome database or custom, as well as TMB, either using the absolute number of mutations, or mutations per expressed MB of genome, using a user-defined threshold. Mutations can also be plotted from a protein point of view using user-defined occurrence cutoffs for showing and annotating mutations. The overall set of 3,217 SCAN-B patients can be filtered based on combinations of clinicopathological and molecular markers (histological type, ER, PgR, HER2, Ki67, NHG, and PAM50 subtype) and treatments (endocrine, chemotherapy, and HER2 treatment). Plots in PDF format as well as the mutation set underlying the currently active plot in tab-separated values (TSV) format can be downloaded for further analysis. The application is based on R Shiny and the source code is available under the BSD 2-clause open source license at http://github.com/cbrueffer/MutationExplorer.

**Figure 8:**
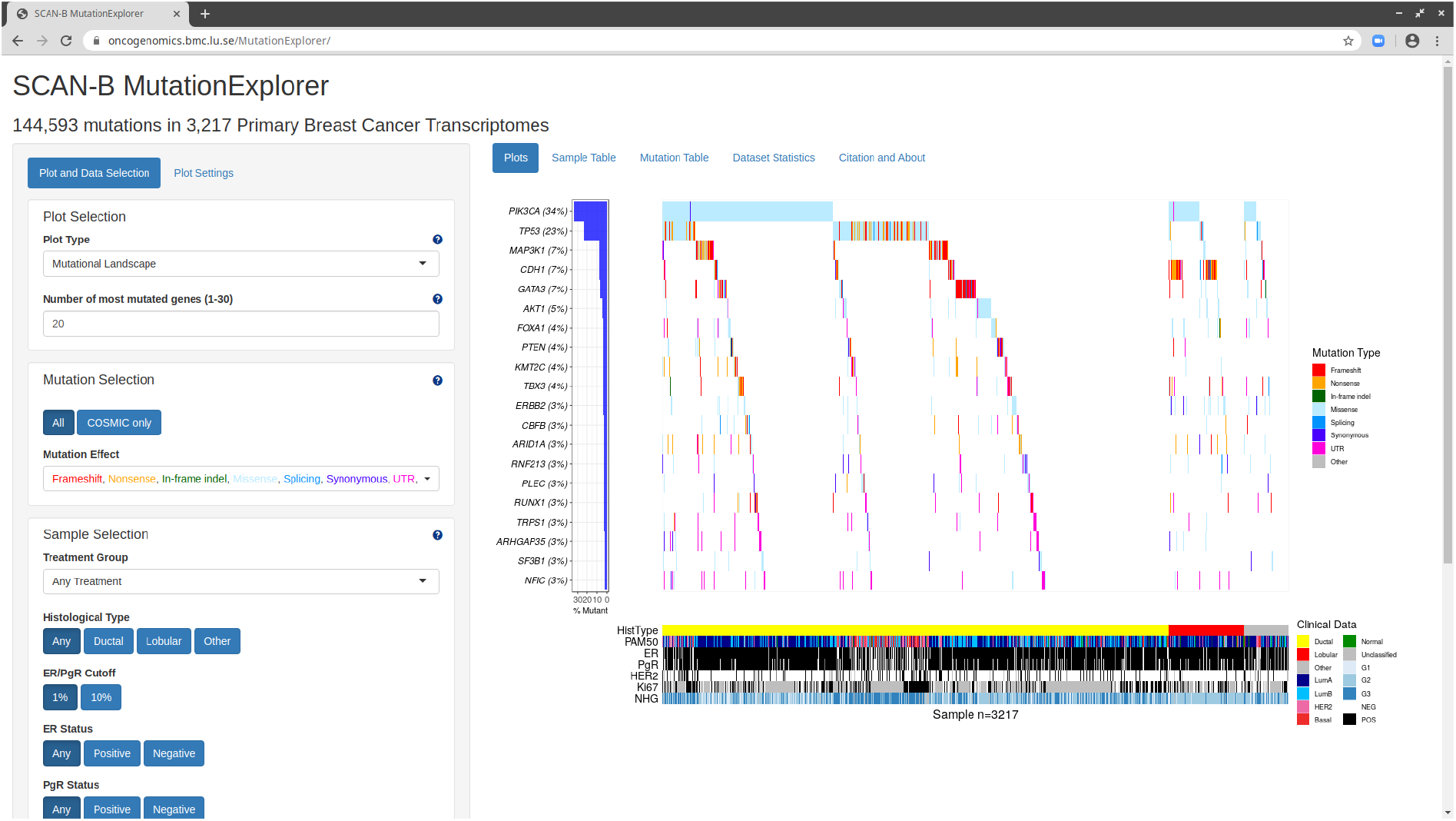
The SCAN-B MutationExplorer web-based application for interactive exploration of mutations, and their association with clinicopathological subgroups and overall survival.

## Discussion

Tumor somatic mutation status is a crucial piece of information for the future of precision medicine to guide treatment selection, and give insight into tumor evolution. For example the PI3K inhibitor alpelisib (Novartis) was recently FDA-approved for use in *PIK3CA*-mutated metastatic BC [64], and the anti-PD1 immunotherapy pembrolizumab was the first drug approved across multiple tumor types based on biomarker status alone [65]. Analysis of DNA is the gold standard for detecting SNVs, indels and larger structural variants. However, many interesting tumor properties are only accessible on the transcriptome level and cannot be interrogated using DNA; most prominently gene expression at the isoform and gene level, as well as *de novo* transcripts originating from gene fusions. The SCAN-B initiative [13] decided early on to perform RNA-seq on the tumors of all enrolled patients. Based on this we have developed, refined, and benchmarked gene expression signatures [10, 66, 67, 68, 69], and detected recurring fusions affecting miRNAs [70]. Detection of somatic mutations from RNA-seq data adds another layer to information that can now be obtained from a single sequencing analysis within one week of surgery [13]. Mutation calling from RNA-seq is not as well refined as that from DNA, although calling from DNA cannot be considered a solved problem given that false positive calls are still prevalent and significant discordance between mutation detection algorithms exists [71, 72, 73]. Compared to DNA, RNA-seq mutation calling is complicated by several factors: additional steps in the library preparation protocols [74, 75], the inherent complexity of the transcriptome resulting in noise, and more complex read alignment accounting for splice sites. This results in potential false positive variant calls due to cDNA synthesis, RNA editing (e.g., `A>I` editing resulting in `A>G` variant calls) [76] and other RNA modifications, as well as read mis-alignment. An inherent limitation is that only mutations in sufficiently expressed regions can be detected.

To date several approaches for RNA-seq mutation calling, mostly in combination with matched DNA, have been developed [77, 78, 79, 80, 81, 82], however calling from RNA-seq alone, particularly from tumor-only samples, is still a challenge. With the advance of targeted and whole exome sequencing into the clinics, and efforts such as TCGA, MSK-Impact, and others, variant calling from DNA-seq has improved in recent years. Part of this improvement is the availability of validation resources such as the Genome in a Bottle datasets [17]. With clinical interest in RNA-seq only recently picking up, e.g. as shown by two recent review articles [11, 12], comparably well-characterized RNA-seq datasets for validation do not yet exist to our knowledge.

The strategy for mutation calling herein was to perform initial variant calling with low requirements on coverage and base quality in order to increase sensitivity while allowing false-positives. To increase specificity we then applied stringent post-hoc filtering that can be easily amended as further annotation data becomes available, or as existing sources receive updates. The advantage of this two-step strategy is the possibility to accommodate different research and clinical questions in the future that may have different filtering needs.

Two major contributors of false-positive mutation calls are germline SNPs/indels and RNA editing. Common approaches for dealing with germline events are calling mutations from matched tumor/normal samples, or filtering SNPs present in databases such as dbSNP. The latter is problematic, since some dbSNP entries with a low variant allele frequency (VAF) may be legitimate somatic mutations that should not be filtered away. On the other hand, filtering on the dbSNP “common” flag (at least 1% VAF in any of the 1000 genomes populations) can lead to many low-VAF germline SNPs remaining. We tried to address this issue by combining the dbSNP and COSMIC databases, and only filtering variants present in dbSNP if they were not present in COSMIC. We filtered out known RNA editing sites using publicly available databases, however there is still an overabundance of `T>C` substitutions in our RNA-based calls compared to DNA-based calls, suggesting many unknown editing sites and insufficient filtering (Figure 3B). Approaches have been developed to identify RNA editing sites using DNA/RNA-trained machine learning models [39] or RNA-seq data alone [83], which may provide ways to improve filtering in the future by creating a SCAN-B RNA editing database.

The two most frequently mutated genes in our study were *PIK3CA* (34% of samples) and *TP53* (23%) in line with previous studies such as TCGA [1], followed by other known drivers *MAP3K1* (7%), *CDH1* (7%), *GATA3* (7%), and *AKT1* (5%) (Figure 2). Other frequently mutated genes in our set such as *RNF213* have been implicated in breast cancer, and for *PLEC* mutations have been reported [78, 84]. While mutation frequencies in oncogenes such as *PIK3CA* are generally in line with previous studies, frequencies in tumor suppressor genes were generally lower in RNA-seq than would be expected from our study population. For example our *TP53* RNA-seq somatic mutation frequency of 23% (reference: 36%, cBioPortal.org) suggests we may be missing a significant fraction of *TP53* mutations present in DNA. Similar trends can be seen in *PTEN* (observed: 3.6%, reference: 4.6%), *BRCA1* (observed: 0.2%, reference: 1.6%), *BRCA2* (observed: 0.03%, reference: 2.2%). This is not surprising since only mutations in sufficiently highly expressed genomic regions can be detected by RNA-seq, and loss of expression of tumor suppressor genes is a hallmark of oncogenesis. Furthermore, truncated mRNAs caused by nonsense mutations are typically removed by nonsense-mediated decay before they can be captured for sequencing. Thus, our findings do not reflect the true mutational spectrum of tumor suppressor genes. Despite these limitations, we could identify a putative mutation in at least one gene targeted by an existing drug in the majority of patient tumors (86.8%), demonstrating that it should be feasible to match most patients to targeted treatments using RNA-seq analyses.

PI3K/AKT/mTOR is a key oncogenic pathway in BC that is frequently activated by alterations in genes such as *PIK3CA*, *MAP3K1*, *AKT1*, or *PTEN*, leading to increased growth signaling. Missense mutations in the *PIK3CA* hotspot residues 1047, 545, and 542 (Figure 3D) have been shown to lead to constituent pathway activation [85]. In a pooled analysis of *PIK3CA* mutation impact on different survival endpoints Zardavas *et al* [60] showed that *PIK3CA* mutant tumors were associated with slightly better 5-year OS than *PIK3CA*-wt tumors in univariable analysis (hazard ratio, 0.89), but not when correcting for clinicopathological and treatment variables. This survival benefit is not reflected in our data, where the only genotype-dependent survival difference for *PIK3CA* is within the TNBC patient group (P=0.049, Figure S4). This difference could be an effect of the treatment regimes, study population, and/or population-based nature of our cohort. Various agents targeting PI3K are in clinical trials, for which *PIK3CA* mutations are predictive biomarkers. Importantly the drug alpelisib (Novartis) [86] has been shown to confer beneficial progression-free survival in HR+/HER2-*PIK3CA* mutant tumors in combination with fulvestrant [64]. This survival benefit may be dependent on co-mutation status of *PIK3CA* and *MAP3K1*; the latter is a tumor suppressor gene in the PI3K/AKT/mTOR pathway whose loss of expression activates the pathway and desensitizes the tumor to PI3K inhibition [87]. In fact we frequently observed inactivating (frameshift/nonsense) *MAP3K1* alterations in *PIK3CA* mutant tumors (77/1,095, 7%). *AKT1* is a known oncogene in this pathway, and had one of the most mutated hotspots (E17K) in the cohort. This mutation leads to constitutive AKT1 signaling and PI3K pathway activation, and sensitizes tumors to the ATP-competitive AKT1 inhibitor AZD5363 (AstraZeneca), but not allosteric inhibitors [54, 88]. Yi *et al* suggest that other, rarer, mutations in *AKT1* are non-sensitizing passenger mutations [88].

Another potential biomarker for PI3K inhibitor efficacy is *PTEN*, the key negative regulator of the PI3K/AKT/mTOR pathway. PTEN protein expression is frequently lost in BC, e.g. caused by gross rearrangements of the gene [61]. The hotspot mutations we identified are of unknown significance (Figure 3E). *PTEN* mutation status alone only significantly stratified patients receiving no systemic treatment into OS low/high risk groups, however when combined with low gene expression in the *PTEN*-MutExp low group mutational information contributed to better stratification (Supplementary Figure S3) than achieved by *PTEN* expression alone, particularly in patients receiving endocrine treatment only (Figure 5), and the HR+/HER2-subgroup (Supplementary Figure S4). This suggests that SNVs and indels are a minor mechanism of PTEN loss in early BC compared to larger deletions, structural rearrangements, and other means of *PTEN* expression loss. Taken together, we detected mutations in multiple PI3K/AKT/mTOR signaling nodes (1,872/3,217 [58.2%] mutated samples) that lead to increased pathway activation and have emerging clinical utility in luminal BC, e.g. through combination with EGFR inhibition as demonstrated in basal-like BC [89].

Loss of p53 activity, either through loss of function (LoF) mutations, dominant-negative mutations, or low expression, is a major contributor to tumorigenesis. While we generally underdetect *TP53* mutations, the identified hotspot residues remain the same as reported in the IARC TP53 database. The vast majority of detected mutations are located in the DNA-binding domain, and 77.6% of overall mutations are missense mutations, likely leading to protein LoF. *TP53* mutations correlated with triple-negative biomarker status (P<0.001) and basal-like PAM50 subtype (P<0.001), as reported before [1]. Clinically these mutations could already be actionable, as *TP53* mutations are a sign of DNA damage-repair deficiency and may be prognostic for sensitivity to PARP inhibition [90, 91]. Recent simulations suggest that a subset of mutations could lead to the formation of a druggable binding pocket, potentially opening therapeutic options to restore p53 function [92]. *TP53* mutant patients had significantly worse OS in the patient subgroups treated only with endocrine therapy, or no systemic treatment at all (Figure 5). This finding was corroborated by worse survival of *TP53* mutant patients in the HR+/HER2-clinical subgroup (Supplementary Figure S4), suggesting that *TP53* mutations identify a subgroup of patients that are spared chemotherapy or systemic therapy overall by appearing low-risk, but are in fact high-risk patients that should be treated accordingly.

ER receptor protein expression is arguably the most important biomarker in BC, with 2,786 (86.6%) tumors of our cohort being ER+. ER positivity is a strong prognostic and treatment predictive biomarker for endocrine treatment efficacy. Several tumors in our cohort harbored *ESR1* mutations, including known endocrine treatment resistance mutations, that are discussed elsewhere in detail (Dahlgren *et al*, submitted). Transcription factors encoded by *GATA3* and *FOXA1* are directly involved in modulating *ESR1* signaling, and their expression is independently associated with beneficial survival in ER+ tumors [93]. While the role of mutations in these genes has not been thoroughly characterized, Takaku *et al* suggest that *GATA3* can function as either oncogene or tumor suppressor depending on the mutations the gene accumulated, and which part of the protein product is impacted [94]. According to their classification the most frequent mutation in our cohort, the P409fs frameshift mutation, results in an elongated protein product compared to *GATA3*-wt that has favourable survival compared to mutations of the second Zinc finger domain.

*ERBB2* (*HER2*) mutations have emerged as a novel biomarker and occur by the majority in patients without *HER2* amplification [2], but also in HER2-amplified cases [53]. Evidence is mounting that recurrent *ERBB2* mutations lead to increased activation of the HER2 receptor in tumors classified as HER2-normal [2, 51, 95]. Activating *ERBB2* mutations have been shown to confer therapy resistance against standard of care drugs such as trastuzumab and lapatinib [53], but can be overcome using pan-HER tyrosine kinase inhibitors such as neratinib [2, 96, 97, 53]. *ERBB2* mutations have also been shown to confer resistance to endocrine therapy in the metastatic setting [98], where HER2-directed drugs are effective [99]. In our study patients with *ERBB2* mutant tumors had markedly worse prognosis than *ERBB2*-wt patients, particularly in ER+/HER2-BC (Figure 5C, Supplementary Figure S4C). Detecting these patients early could open up additional treatment options to patients, especially when they show signs of resistance to endocrine therapy. Our results also demonstrate that mutation and amplification can already co-occur in early, treatment-naïve BC. In the metastatic setting Neratinib monotherapy has been found effective in these patients, suggesting it may be a useful part of the treatment arsenal for early BC as well.

The role of alternative splicing in tumorigenesis has recently garnered increased attention, and the extend of isoform switching in several cancer types, including BC, has been characterized [100]. *SF3B1* encodes a subunit of the spliceosome and mutations in this gene have been identified as potentially interesting treatment targets after having been observed in myelodysplastic syndromes and chronic lymphocytic leukemia. The K700E hotspot mutation deregulates splicing and results in differential splicing patterns in BC [55]. The clinical effect of these mutations is unclear, however we did not detect significant survival stratification in important biomarker or treatment groups.

Mutations in *CBFB* were recently identified as recurrent in ER+/HER2-BC [57]. CBFB is a transcriptional co-activator of Runx2, an expression regulator of several genes involved in metastatic processes such as cell migration. Increased CBFB expression has been identified as essential for cell invasion in BC [101]. While we could confirm occurrence and mutation frequency in ER+ BC, we did not observe the splice site mutation described by Griffith *et al* [57], perhaps due to degradation of the spliced mRNA by nonsense-mediated decay.

### Pathways

Individual mutations, particularly in infrequently mutated genes, affect a smaller number of molecular pathways to achieve the classical hallmarks of cancer such as sustained proliferative signaling. The two most eminent co-mutation clusters we observed in a subset of tumors enriched for basal-like and HER2-enriched molecular subtypes were one made up of hedgehog signaling, p53-independent DNA repair, and hypoxia response, the other of co-mutated NOTCH1/2/3 signaling pathways (Figure 4). Both clusters are linked in their relation to cancer stem cell development [102, 103], which, in addition to the NOTCH and Hedgehog pathways themselves, has emerged as a novel treatment target, particularly in TNBC. A dominant cluster of luminal tumors was co-mutated in the PI3K/AKT, MET, RET, EGFR, ERBB2, and ERBB4 signaling pathways. Activation of these pathways is involved in the development of ER+ BC through proliferation-inducing signaling, or endocrine therapy resistance, e.g. via activating *ERBB2* mutations [98].

Mutation status of several individual pathways was associated with reduced OS in different treatment and clinical biomarker subgroups. In patients not systemically treated or only treated with endocrine therapy WNT, NOTCH2, p53-independent DNA repair pathway mutation status, and Hedgehog signaling mutation status may identify patients diagnosed as low risk who may benefit from more adjuvant treatment (Figure 6). While these stratification profiles were visible in treatment subgroups, they mostly did not yield significant results in clinical biomarker subgroups (Supplementary Figure 6). This may indicate that current risk stratification in histopathological biomarker subgroups is inadequate and should take molecular information into account – something we and others have also shown on the level of gene expression [10].

Identifying the mutation status of pathways and pathway clusters may aid in future clinical trials and treatment, e.g. by aiding selection of treatments that exploit synthetic lethality [104].

### Tumor Mutational Burden

High TMB has been identified as a predictive biomarker for response to immune checkpoint therapy in diverse solid tumors [63, 105, 106, 107, 108]. Using RNA-seq to assess mutational burden may be a useful capability for clinical trials and eventual clinical implementation in BC [109]. Questions remain however, as TMB is influenced by many biological and technical factors such as ploidy, tumor heterogeneity and clonality (ASCO SITC 2019, Abstract 27), sample tumor cell content, sequencing depth, and variant filtering. Which cutoff to use for stratifying patients into TMB groups is also still emerging [109, 110], and specifically in RNA-seq has to our knowledge not been addressed. Due to this, and to account for different expression profiles per tumor, we decided to use the median amount of non-synonymous mutations per MB of transcriptome across the cohort (0.08 mutations) to stratify patients into TMB high and low groups and use it to study OS in different conventional treatment and biomarker subgroups. In several of these groups high TMB was significantly associated with worse survival, confirming previous reports [111], however interestingly not in TNBC. These tumors typically show higher TMB than other clinical BC subtypes, likely because many of them have impaired DNA damage repair mechanisms. Shah *et al* [112] showed that only approximately 36% of mutations in TNBCs are expressed; we speculate that due to this we may underestimate TMB in several of our TMB-low patients. Additionally, RNA-seq underdetects truncating mutations such as frame-shift indels that are a major source of neoantigens. Immune checkpoint therapy is a particularly attractive treatment approach in patients with TNBC and basal-like tumors for which currently no targeted therapy exists. For these patients, determination of TMB using DNA-seq may be a better option than relying on RNA-seq.

### SCAN-B MutationExplorer

Large scale projects such as TCGA have generated vast amounts of data, but bioinformatics skills are required to make efficient use of them. Web portals such as cBioPortal [113] have emerged to make these huge datasets explorable without specialized skills. In this spirit we developed the open source web application SCAN-B MutationExplorer to make our mutation dataset easily accessible for other researchers. We hope that SCAN-B MutationExplorer will aid knowledge generation and the development of better BC biomarkers in the future. The open source nature of the portal allows developers to adopt the code for their own purposes, and we welcome contributions of any kind.

### Limitations

The mutation calling we have performed herein tries to achieve sensitive variant calling by using lenient parameters, and heavy filtering of the resulting variants based on stringent quality factors, annotations, and curated databases. This approach has several limitations. While our 275 patient cohort for filter development had matched tumor and normal DNA sequencing data, the SCAN-B cohort only consisted of tumor RNA-seq data. This made accounting for PCR and sequencing artifacts more challenging. Further while many germline events can be filtered by comparing to general databases such as dbSNP, and population-specific ones such as SweGen, these databases are incomplete and it is thus not possible to remove all germline events this way. As these databases improve, our filters can be upgraded to increase performance. Herein we also applied filters developed in a matched DNA/RNA set of targeted capture sequencing of 1,697 genes and 1,047 miRNAs (275 sample ABiM cohort) to whole mRNA-seq (3,217 sample SCAN-B cohort). This assumes the transcriptional characteristics of the captured regions are representative for the whole mRNA.

## Conclusions

We presented a tumor-only RNA-seq variant calling strategy and resulting mutation dataset from a large population-based early breast cancer cohort. Although variant calling from RNA-seq data is limited to expressed regions of the genome, mutations in important BC genes such as *PIK3CA*, *TP53*, and *ERBB2*, as well as pathways can be reliably detected, allowing to inform clinical trials and eventual reporting to the clinic. Mutations in *TP53*, *PIK3CA*, *ERBB2*, and *PTEN* provided prognostic information in several treatment and biomarker patient subgroups, demonstrating the utility of the dataset for research. We make this dataset available for analysis and download via the open source web application SCAN-B MutationExplorer, accessible at http://oncogenomics.bmc.lu.se/MutationExplorer.

## Data Availability

Clinical data for all samples, as well as mutation calls can be downloaded from the SCAN-B MutationExplorer website. For access to the full dataset, please contact the corresponding author. Gene expression data in FPKM is available from the NCBI Gene Expression Omnibus (accession number GSE96058).

## Author Contributions

CB, CW, and LHS conceived the study. CB, SG, CW, JV-C, JH, AMG, YC, NL, and LHS analyzed data. JV-C, CH, JH, AE, CL, NL, MM, LR, ÅB, and LHS established the SCAN-B initiative. CB, CW, and LHS established the DNA-seq analyses. CB, SG, and LHS established the RNA-seq mutation calling pipeline and filters. JV-C, CH, JH, AE, CL, NL, MM, LR, ÅB, and LHS provided clinical information. LHS supervised the project, and CB and LHS wrote the report with assistance from all authors. All authors discussed, critically revised, and approved the final version of the report for publication.

## Acknowledgements

We thank the patients who were part of this study and the SCAN-B study, the employees of the SCAN-B laboratory, the South Sweden Breast Cancer Group, and all SCAN-B collaborators at Hallands Hospital Halmstad, Helsingborg Hospital, Blekinge County Hospital, Central Hospital Kristianstad, Skåne University Hospital Lund/Malmö, Central Hospital Växjö, for inclusion of patients and sampling of tissue for this study. We also thank the Swedish National Breast Cancer Registry and Regional Cancer Center South for clinical data. The SweGen allele frequency data was generated by Science for Life Laboratory.

This work was supported by the Mrs. Berta Kamprad Foundation and funded in part by the Swedish Research Council, Swedish Cancer Society, Swedish Foundation for Strategic Research, Knut and Alice Wallenberg Foundation, VINNOVA, Governmental Funding of Clinical Research within National Health Service (ALF), Scientific Committee of Blekinge County Council, Crafoord Foundation, Lund University Medical Faculty, Gunnar Nilsson Cancer Foundation, Skåne University Hospital Foundation, BioCARE Research Program, King Gustav Vth Jubilee Foundation, and the Krapperup Foundation.

## Conflict of Interest

CB, SG, AMG, YC, and LHS are shareholders and/or employees of SAGA Diagnostics AB. LHS has received honorarium from Novartis and Boehringer-Ingelheim. All remaining authors have declared no conflicts of interest.

---

**Figure S1:**
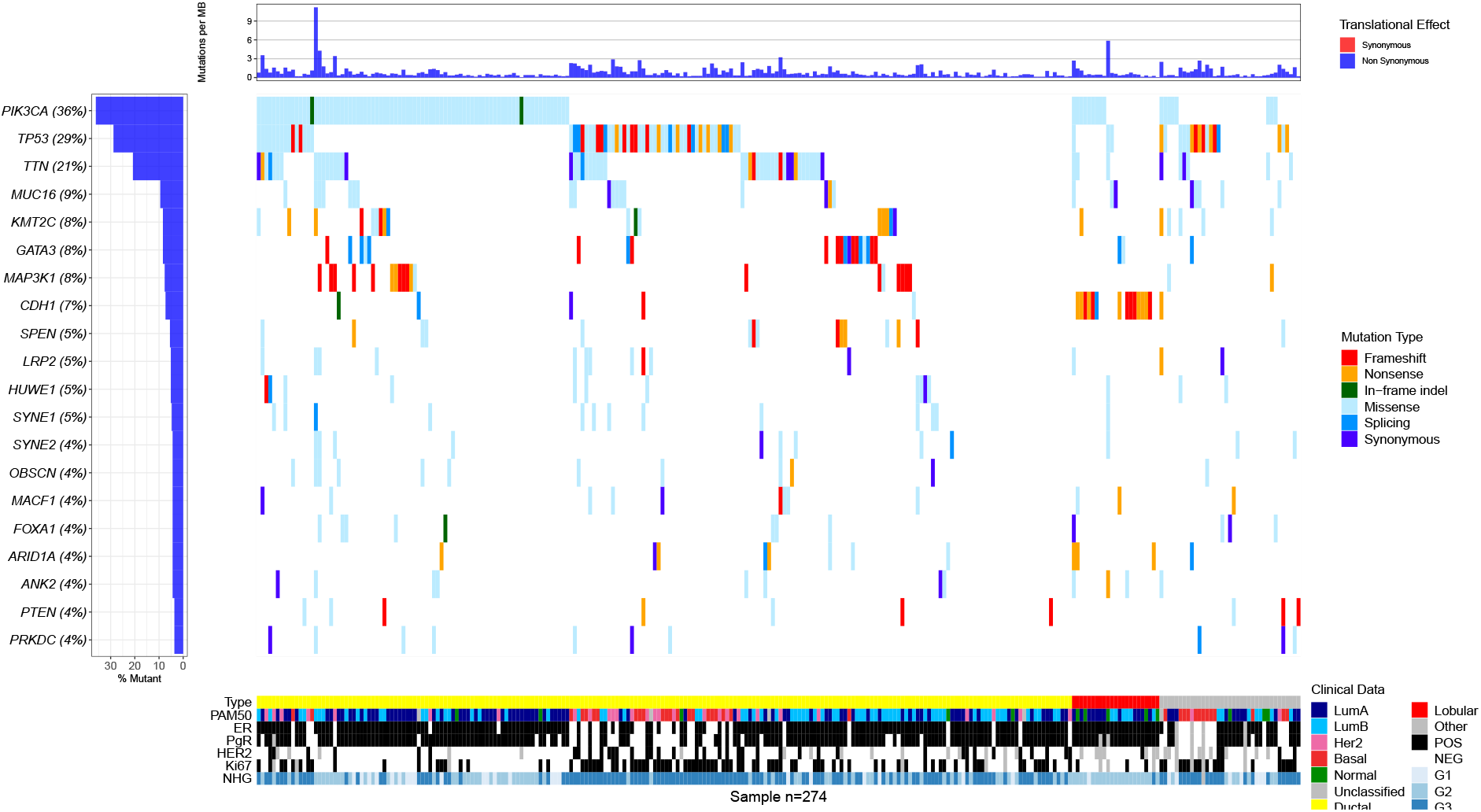
Waterfall plot of the 20 most mutated genes in the targeted DNA-seq ABiM cohort data.

**Figure S2:**
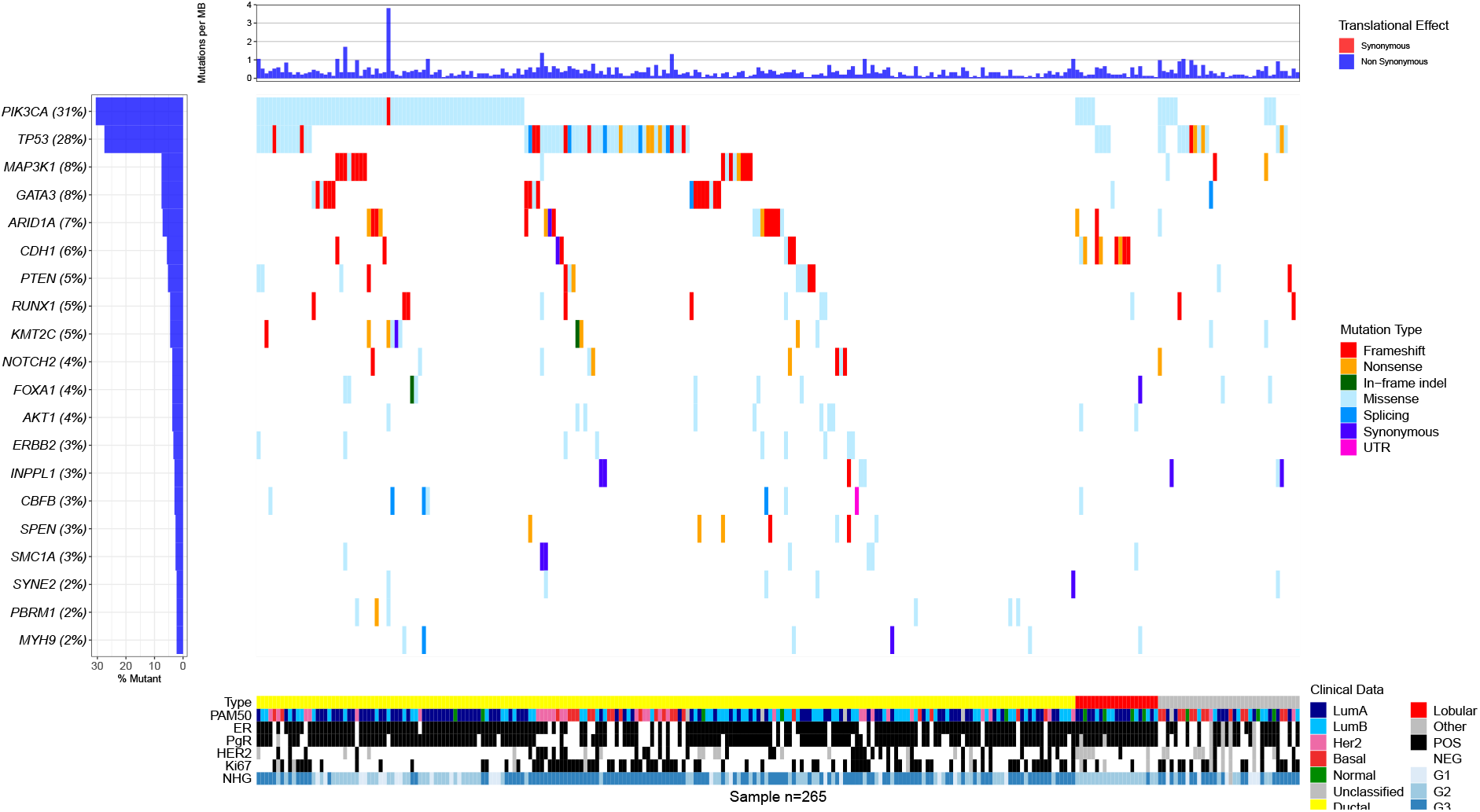
Waterfall plot of the 20 most mutated genes in RNA-seq ABiM cohort data.

**Figure S3:**
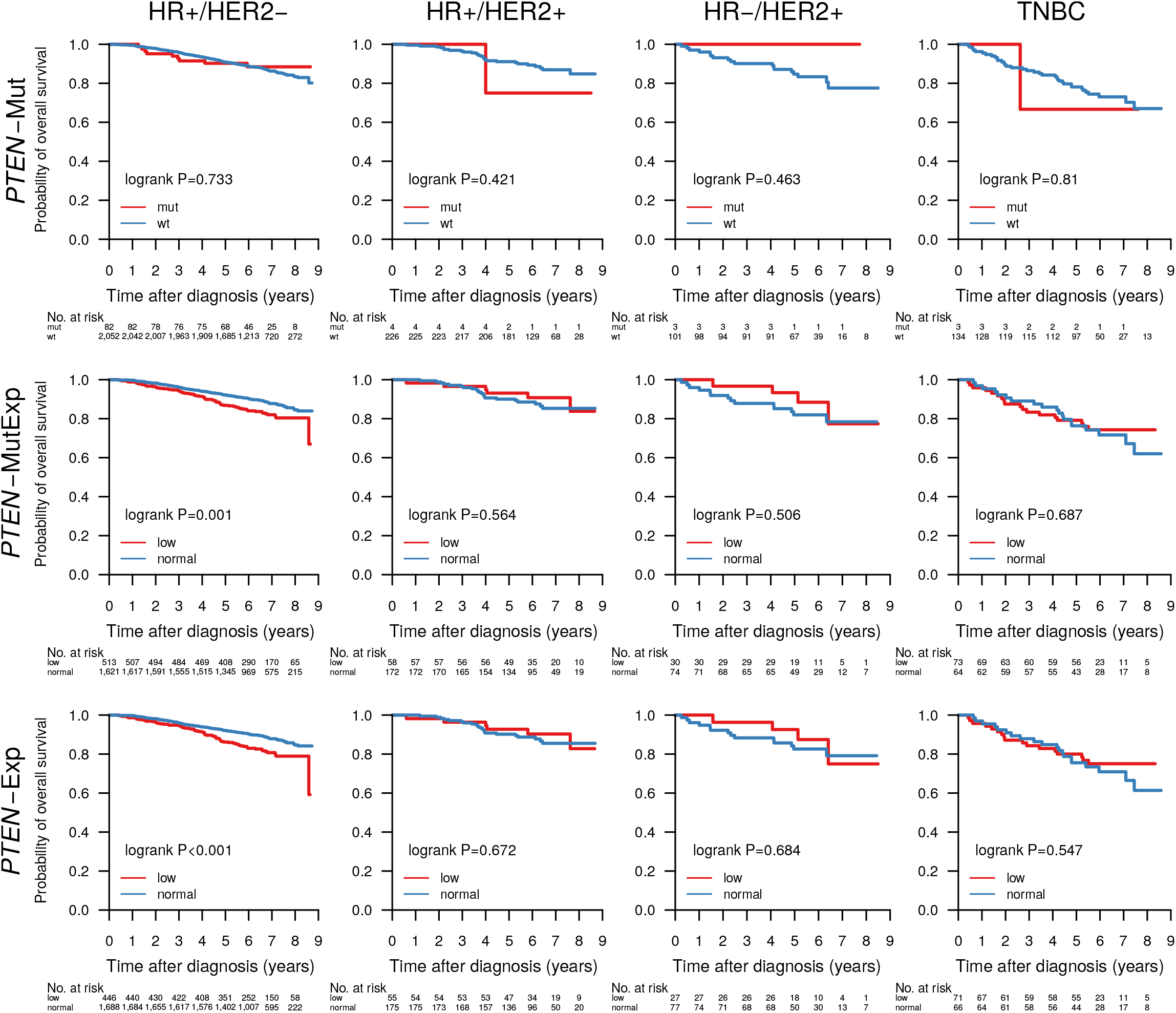
Overall survival of patients with tumors containing a *PTEN* mutation (*PTEN*-Mut), a *PTEN* mutation and/or *PTEN* expression in the lower cohort-quartile (*PTEN*-MutExp), or *PTEN* expression in the lower cohort-quartile (*PTEN*-Exp) in the clinical patient subgroups HR+/HER2-(HR+ when ER+ and PgR+, HR-otherwise), HR+/HER2+, HR-/HER2+, and TNBC.

**Figure S4:**
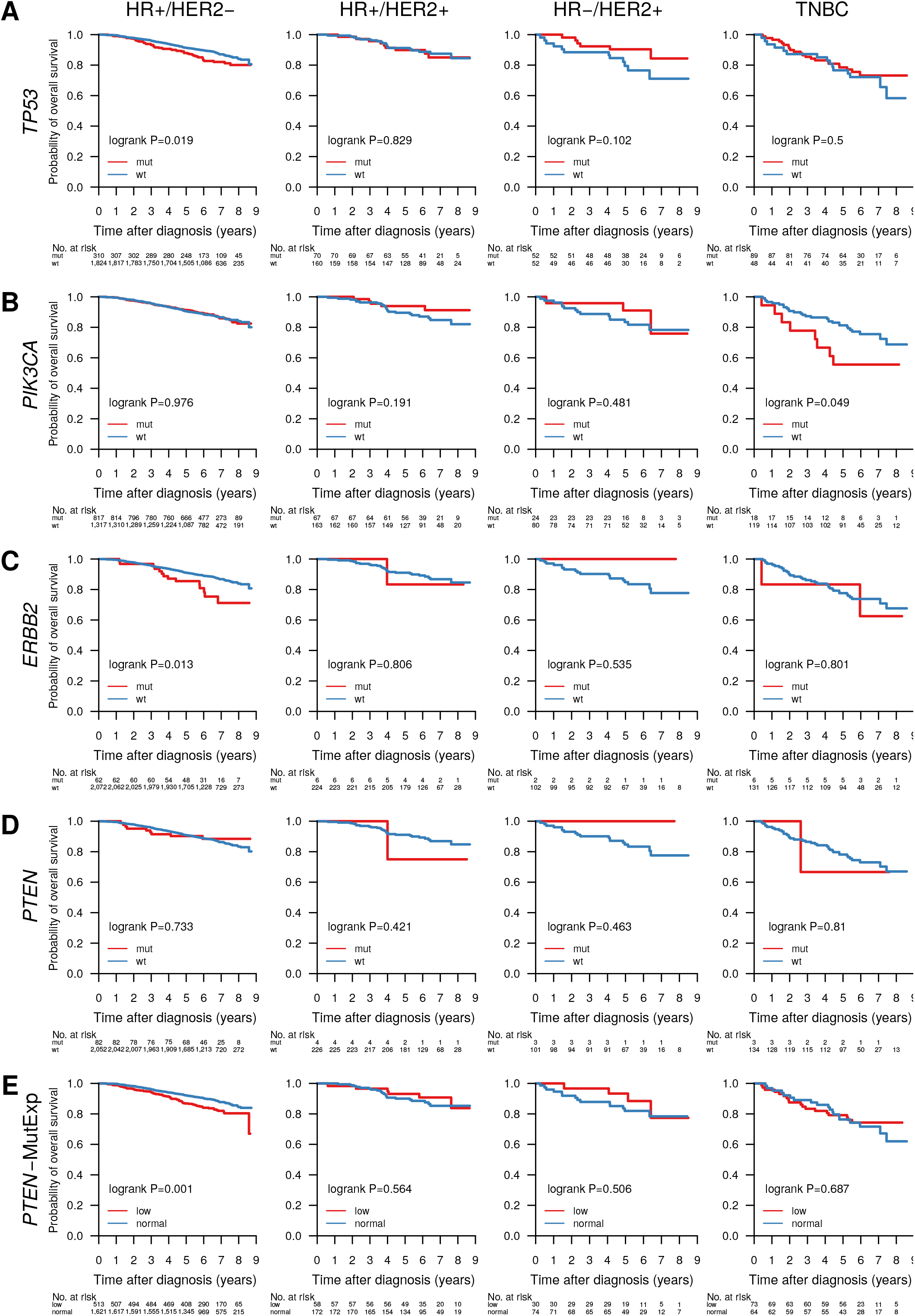
Overall survival of patients with tumors containing mutations in the genes **A**) *TP53*, **B**) *PIK3CA*, **C**) *ERBB2*, **D**) *PTEN*, **E**) *PTEN* or *PTEN* expression in the lower quartile across the cohort, stratified by the clinical patient subgroups HR+/HER2-(HR+ when ER+ and PgR+, HR-otherwise), HR+/HER2+, HR-/HER2+, and TNBC.

**Figure S5:**
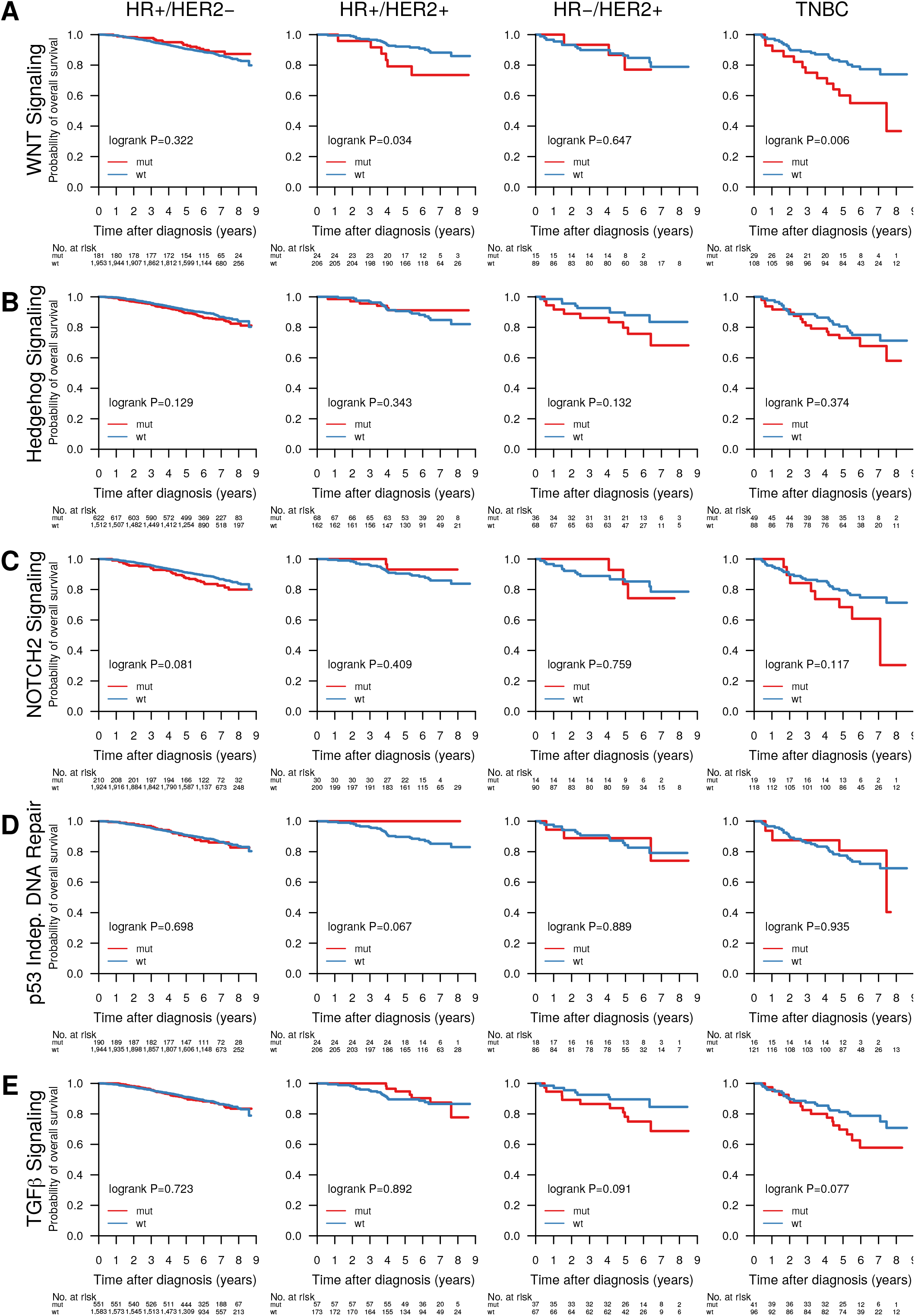
Overall survival of patients with tumors containing mutations in pathways WNT signaling, Hedgehog signaling, cell cycle, p53 independent DNA damage repair, and TGF*β* signaling, stratified by the clinical patient subgroups HR+/HER2-(HR+ when ER+ and PgR+, HR-otherwise), HR+/HER2+, HR-/HER2+, and TNBC.

**Figure S6:**
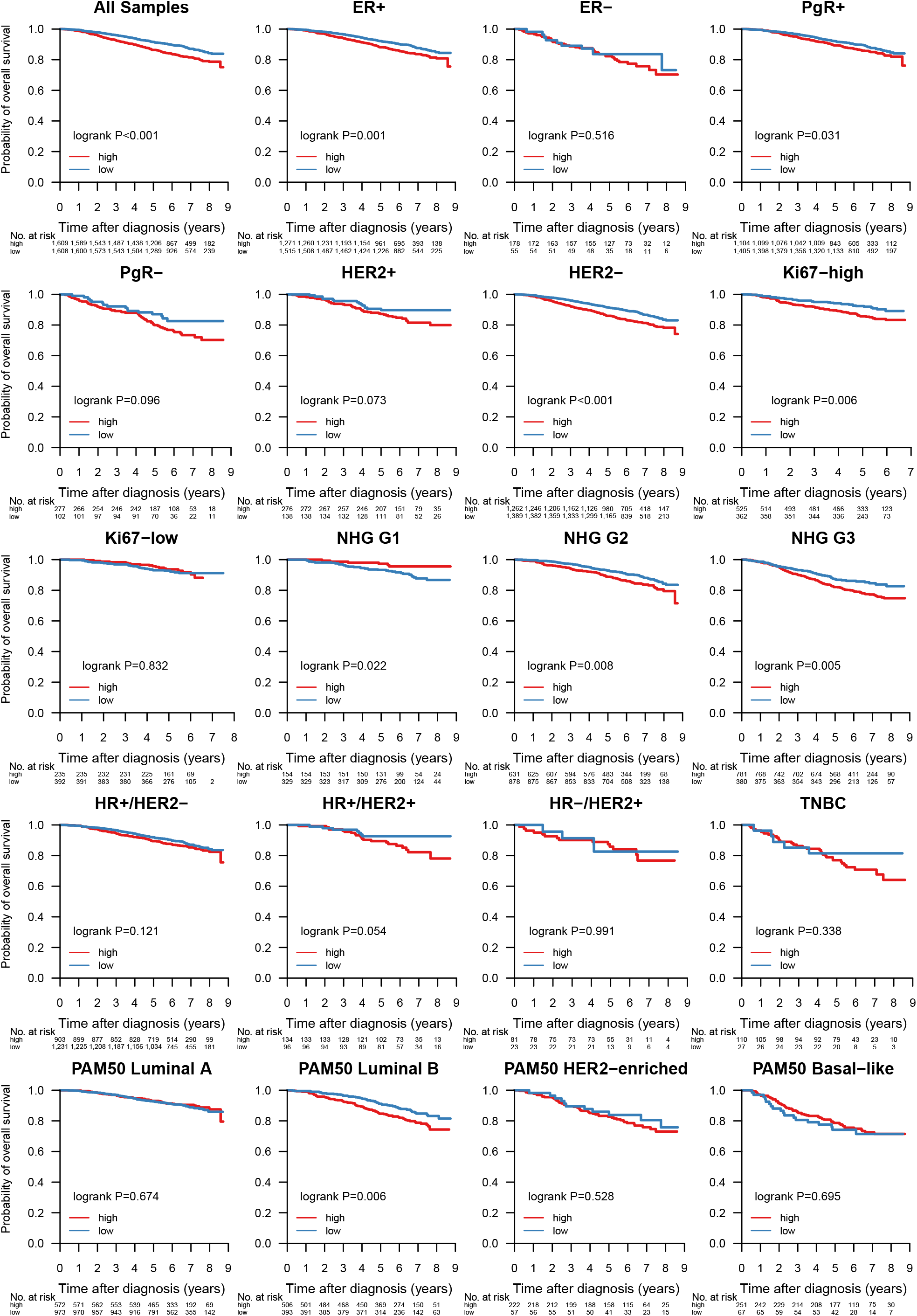
Association of TMB with OS in all samples, and within the biomarker patient subgroups ER+, ER-, PgR+, PgR-, HER2 amplified, HER2 normal, Ki67-high, Ki67-low, NHG 1-3, HR+/HER2-, HR+/HER2+, HR-/HER2+, TNBC, and PAM50 Luminal A and B, HER2-enriched, and basal-like.

**Table 1:** Patient demographics and clinicopathological variables in the ABiM and SCAN-B cohorts.

**Table 2:** Number of Mutations in the ABiM (DNA-seq and RNA-seq) and SCAN-B (RNA-seq) cohorts. Sample numbers differ from total cohort sizes due to filtering resulting in samples with no remaining post-filter mutations.

**Table 3:** The most occurring non-synonymous mutations in the genes *PIK3CA*, *AKT1*, *SF3B1*, *GATA3*, *ERBB2*, *TP53*, *FOXA1*, and *CDH1* in 3,217 SCAN-B samples. Shown are the total number of mutations, the frequency of the mutations in the SCAN-B cohort (Mut. Samples), and the frequency of a particular mutation within all mutations in the gene (Mut in Gene).

**Table 4:** One-sided Fisher’s exact test P-values for association of gene mutation status and clinical/molecular biomarker status. Associations with P-values below the significance threshold of 0.05 are highlighted in green.

**Supplementary Table S1:** RNA-seq statistics for the ABiM and SCAN-B cohorts.

**Supplementary Table S2:** RNA-seq variant filters and descriptions.

**Supplementary Table S3:** Number of alterations by type in the post-filter SCAN-B RNA-seq mutation set stratified by single nucleotide variants (SNVs), insertions, and deletions.

**Supplementary Table S4:** Reactome pathway name to pathway ID mapping.

## Notes

http://oncogenomics.bmc.lu.se/MutationExplorer

## References

[1] The Cancer Genome Atlas. Comprehensive molecular portraits of human breast tumours. Nature. 2012;490(7418):61–70.

[2] Bose R, Kavuri SM, Searleman AC, Shen W, Shen D, Koboldt DC, et al. Activating HER2 mutations in HER2 gene amplification negative breast cancer. Cancer Discovery. 2013;3(2):224–237.

[3] Robinson DR, Wu YM, Vats P, Su F, Lonigro RJ, Cao X, et al. Activating ESR1 mutations in hormone-resistant metastatic breast cancer. Nature Genetics. 2013;45:1446–1451.

[4] Garcia-Murillas I, Schiavon G, Weigelt B, Ng C, Hrebien S, Cutts RJ, et al. Mutation tracking in circulating tumor DNA predicts relapse in early breast cancer. Science Translational Medicine. 2015;7(302).

[5] Förnvik D, Aaltonen KE, Chen Y, George AM, Brueffer C, Rigo R, et al. Detection of circulating tumor cells and circulating tumor DNA before and after mammographic breast compression in a cohort of breast cancer patients scheduled for neoadjuvant treatment. Breast Cancer Research and Treatment. 2019;177(2):447–455.

[6] Ciriello G, Gatza ML, Beck AH, Wilkerson MD, Rhie SK, Pastore A, et al. Comprehensive Molecular Portraits of Invasive Lobular Breast Cancer. Cell. 2015;163(2):506–519.

[7] Cheng DT, Mitchell TN, Zehir A, Shah RH, Benayed R, Syed A, et al. Memorial Sloan Kettering-integrated mutation profiling of actionable cancer targets (MSK-IMPACT): A hybridization capture-based next-generation sequencing clinical assay for solid tumor molecular oncology. Journal of Molecular Diagnostics. 2015;17(3):251–264.

[8] Ma C, Shao M, Kingsford C. SQUID: Transcriptomic Structural Variation Detection from RNA-seq. Genome Biology. 2018;19(52).

[9] Talevich E, Shain AH. CNVkit-RNA: Copy number inference from RNA-Sequencing data. bioRxiv. 2018;(408534).

[10] Brueffer C, Vallon-Christersson J, Grabau D, Ehinger A, Häkkinen J, Hegardt C, et al. Clinical Value of RNA Sequencing–Based Classifiers for Prediction of the Five Conventional Breast Cancer Biomarkers: A Report From the Population-Based Multicenter Sweden Cancerome Analysis Network—Breast Initiative. JCO Precision Oncology. 2018;(2):1–18.

[11] Byron SA, Van Keuren-Jensen KR, Engelthaler DM, Carpten JD, Craig DW. Translating RNA sequencing into clinical diagnostics: Opportunities and challenges. Nature Reviews Genetics. 2016;17(5):257–271.

[12] Cieślik M, Chinnaiyan AM. Cancer transcriptome profiling at the juncture of clinical translation. Nature Reviews Genetics. 2018;19(2):93–109.

[13] Saal LH, Vallon-Christersson J, Häkkinen J, Hegardt C, Grabau D, Winter C, et al. The Sweden Cancerome Analysis Network - Breast (SCAN-B) Initiative: a large-scale multicenter infrastructure towards implementation of breast cancer genomic analyses in the clinical routine. Genome Medicine. 2015;7(1):1–12.

[14] Rydén L, Loman N, Larsson C, Hegardt C, Vallon-Christersson J, Malmberg M, et al. Minimizing inequality in access to precision medicine in breast cancer by real-time population-based molecular analysis in the SCAN-B initiative. British Journal of Surgery. 2018;105(2):e158–e168.

[15] Sotiriou C, Wirapati P, Loi S, Harris A, Fox S, Smeds J, et al. Gene expression profiling in breast cancer: Understanding the molecular basis of histologic grade to improve prognosis. Journal of the National Cancer Institute. 2006;98(4):262–272.

[16] Roepman P, Horlings HM, Krijgsman O, Kok M, Bueno-de Mesquita JM, Bender R, et al. Microarray-based determination of estrogen receptor, progesterone receptor, and HER2 receptor status in breast cancer. Clinical Cancer Research. 2009;15(22):7003–7011.

[17] Zook JM, Catoe D, McDaniel J, Vang L, Spies N, Sidow A, et al. Extensive sequencing of seven human genomes to characterize benchmark reference materials. Scientific Data. 2016;3.

[18] Li H, Bloom JM, Farjoun Y, Fleharty M, Gauthier LD, Neale B, et al. A synthetic-diploid benchmark for accurate variant-calling evaluation. Nature Methods. 2018;15(8):595–597.

[19] Winter C, Nilsson MP, Olsson E, George AM, Chen Y, Kvist A, et al. Targeted sequencing of BRCA1 and BRCA2 across a large unselected breast cancer cohort suggests one third of mutations are somatic. Annals of Oncology. 2016;27:1532–1538.

[20] Parkhomchuk D, Borodina T, Amstislavskiy V, Banaru M, Hallen L, Krobitsch S, et al. Transcriptome analysis by strand-specific sequencing of complementary DNA. Nucleic acids research. 2009;37(18):e123.

[21] Grüning B, Dale R, Sjödin A, Rowe J, Chapman BA, Tomkins-Tinch CH, et al. Bioconda: sustainable and comprehensive software distribution for the life sciences. Nature Methods. 2018;15(7):475–476.

[22] Tischler G, Leonard S. Biobambam: Tools for read pair collation based algorithms on BAM files. Source Code for Biology and Medicine. 2014;9(1).

[23] Lai Z, Markovets A, Ahdesmaki M, Chapman B, Hofmann O, McEwen R, et al. VarDict: a novel and versatile variant caller for next-generation sequencing in cancer research. Nucleic Acids Research. 2016;44(11):e108.

[24] Zhao H, Sun Z, Wang J, Huang H, Kocher JP, Wang L. CrossMap: A versatile tool for coordinate conversion between genome assemblies. Bioinformatics. 2014;30(7):1006–1007.

[25] Kim D, Langmead B, Salzberg SL. HISAT: a fast spliced aligner with low memory requirements. Nature Methods. 2015;12(4):357–360.

[26] Tarasov A, Vilella AJ, Cuppen E, Nijman IJ, Prins P. Sambamba: Fast processing of NGS alignment formats. Bioinformatics. 2015;31(12):2032–2034.

[27] Faust GG, Hall IM. SAMBLASTER: Fast duplicate marking and structural variant read extraction. Bioinformatics. 2014;30(17):2503–2505.

[28] Köster J, Rahmann S. Snakemake-a scalable bioinformatics workflow engine. Bioinformatics. 2012;28(19):2520–2522.

[29] Pedersen BS, Layer RM, Quinlan AR, Li H, Wang K, Li M, et al. Vcfanno: fast, flexible annotation of genetic variants. Genome Biology. 2016;17(1):118.

[30] Sherry ST, Ward M, Kholodov M, Baker J, Phan L, Smigielski E, et al. dbSNP: the NCBI database of genetic variation. Nucleic Acids Research. 2001;29(1):308–311.

[31] Karczewski KJ, Francioli LC, Tiao G, Cummings BB, Alföldi J, Wang Q, et al. Variation across 141,456 human exomes and genomes reveals the spectrum of loss-of-function intolerance across human protein-coding genes. bioRxiv. 2019;(531210).

[32] Forbes SA, Beare D, Gunasekaran P, Leung K, Bindal N, Boutselakis H, et al. COSMIC: Exploring the world’s knowledge of somatic mutations in human cancer. Nucleic Acids Research. 2015;43(D1):D805–D811.

[33] Sondka Z, Bamford S, Cole CG, Ward SA, Dunham I, Forbes SA. The COSMIC Cancer Gene Census: describing genetic dysfunction across all human cancers. Nature Reviews Cancer. 2018;18(11):696–705.

[34] Griffith M, Spies NC, Krysiak K, McMichael JF, Coffman AC, Danos AM, et al. CIViC is a community knowledgebase for expert crowdsourcing the clinical interpretation of variants in cancer. Nature Genetics. 2017;49(2):170–174.

[35] Ameur A, Dahlberg J, Olason P, Vezzi F, Karlsson R, Martin M, et al. SweGen: A whole-genome data resource of genetic variability in a cross-section of the Swedish population. European Journal of Human Genetics. 2017;25(11):1253–1260.

[36] Maretty L, Jensen JM, Petersen B, Sibbesen JA, Liu S, Villesen P, et al. Sequencing and de novo assembly of 150 genomes from Denmark as a population reference. Nature. 2017;548(7665):87–91.

[37] Ramaswami G, Li JB. RADAR: A rigorously annotated database of A-to-I RNA editing. Nucleic Acids Research. 2014;42(D1).

[38] Kiran A, Baranov PV. DARNED: A DAtabase of RNa editing in humans. Bioinformatics. 2010;26(14):1772–1776.

[39] Sun J, De Marinis Y, Osmark P, Singh P, Bagge A, Valtat B, et al. Discriminative prediction of A-To-I RNA editing events from DNA sequence. PLoS ONE. 2016;11(10):1–18.

[40] Picardi E, Erchia AMD, Giudice CL, Pesole G. REDIportal: a comprehensive database of A-to-I RNA editing events in humans. Nucleic Acids Research. 2017;45(D1):D750–D757.

[41] Gonzalez-Perez A, Perez-llamas C, Deu-Pons J, Tamborero D, Schroeder MP, Jene-Sanz A, et al. IntOGen-mutations identifies cancer drivers across tumor types. Nature Methods. 2013;10(11).

[42] Cotto KC, Griffith OL, Wollam A, Wagner AH, Griffith M, Feng YY, et al. DGIdb 3.0: a redesign and expansion of the drug–gene interaction database. Nucleic Acids Research. 2017;46(D1):D1068–D1073.

[43] Cingolani P, Platts A, Coon M, Nguyen T, Wang L, Land SJ, et al. A program for annotating and predicting the effects of single nucleotide polymorphisms, SnpEff: SNPs in the genome of Drosophila melanogaster strain w1118; iso-2; iso-3. Fly. 2012;6(2):80–92.

[44] Cingolani P, Patel VM, Coon M, Nguyen T, Land SJ, Ruden DM, et al. Using Drosophila melanogaster as a model for genotoxic chemical mutational studies with a new program, SnpSift. Frontiers in Genetics. 2012;3(35).

[45] Skidmore ZL, Wagner AH, Lesurf R, Campbell KM, Kunisaki J, Griffith OL, et al. GenVisR: Genomic Visualizations in R. Bioinformatics. 2016;32(19):3012–3014.

[46] Blokzijl F, Janssen R, Boxtel Rv, Cuppen E. MutationalPatterns: comprehensive genome-wide analysis of mutational processes. Genome Medicine. 2018;10(1).

[47] Pereira B, Chin SF, Rueda OM, Vollan HKM, Provenzano E, Bardwell HA, et al. The somatic mutation profiles of 2,433 breast cancers refines their genomic and transcriptomic landscapes. Nature Communications. 2016;7(11479).

[48] Giacomelli AO, Yang X, Lintner RE, McFarland JM, Duby M, Kim J, et al. Mutational processes shape the landscape of TP53 mutations in human cancer. Nature Genetics. 2018;50(10):1381–1387.

[49] Byrnes G, Ardin M, Bouaoun L, Sonkin D, Olivier M, Zavadil J, et al. TP53 Variations in Human Cancers: New Lessons from the IARC TP53 Database and Genomics Data. Human Mutation. 2016;37(9):865–876.

[50] Saal LH, Johansson P, Holm K, Gruvberger-Saal SK, She QB, Maurer M, et al. Poor prognosis in carcinoma is associated with a gene expression signature of aberrant PTEN tumor suppressor pathway activity. Proceedings of the National Academy of Sciences of the United States of America. 2007;104(18):7564–7569.

[51] Wen W, Chen WS, Xiao N, Bender R, Ghazalpour A, Tan Z, et al. Mutations in the kinase domain of the HER2/ERBB2 gene identified in a wide variety of human cancers. Journal of Molecular Diagnostics. 2015;17(5):487–495.

[52] Ross JS, Gay LM, Wang K, Ali SM, Chumsri S, Elvin JA, et al. Nonamplification ERBB2 genomic alterations in 5605 cases of recurrent and metastatic breast cancer: An emerging opportunity for anti-HER2 targeted therapies. Cancer. 2016;122(17):2654–2662.

[53] Cocco E, Carmona FJ, Razavi P, Won HH, Cai Y, Rossi V, et al. Neratinib is effective in breast tumors bearing both amplification and mutation of ERBB2 (HER2). Science Signaling. 2018;11(551).

[54] Hyman DM, Smyth LM, Donoghue MTA, Westin SN, Bedard PL, Emma J, et al. AKT Inhibition in Solid Tumors With AKT1 Mutations. Journal of Clinical Oncology. 2017;35(20).

[55] Maguire SL, Leonidou A, Wai P, Marchiò C, Ng CK, Sapino A, et al. SF3B1 mutations constitute a novel therapeutic target in breast cancer. The Journal of Pathology. 2015;235(4):571–580.

[56] Alsafadi S, Houy A, Battistella A, Popova T, Wassef M, Henry E, et al. Cancer-associated SF3B1 mutations affect alternative splicing by promoting alternative branchpoint usage. Nature Communications. 2016;7:1–12.

[57] Griffith OL, Spies NC, Anurag M, Griffith M, Luo J, Tu D, et al. The prognostic effects of somatic mutations in ER-positive breast cancer. Nature Communications. 2018;9(1).

[58] Fabregat A, Jupe S, Matthews L, Sidiropoulos K, Gillespie M, Garapati P, et al. The Reactome Pathway Knowledgebase. Nucleic Acids Research. 2018;46(D1):D649–D655.

[59] Jassal B, Matthews L, Viteri G, Gong C, Lorente P, Fabregat A, et al. The reactome pathway knowledgebase. Nucleic acids research. 2020;48(D1):D498–D503.

[60] Zardavas D, te Marvelde L, Milne RL, Fumagalli D, Fountzilas G, Kotoula V, et al. Tumor PIK3CA Genotype and Prognosis in Early-Stage Breast Cancer: A Pooled Analysis of Individual Patient Data. Journal of Clinical Oncology. 2018;36(10):981–990.

[61] Saal LH, Gruvberger-Saal SK, Persson C, Lövgren K, Jumppanen M, Staaf J, et al. Recurrent gross mutations of the PTEN tumor suppressor gene in breast cancers with deficient DSB repair. Nature Genetics. 2008;40(1):102–107.

[62] Zhang HY, Liang F, Jia ZL, Song ST, Jiang ZF. PTEN mutation, methylation and expression in breast cancer patients. Oncology Letters. 2013;6(1):161–168.

[63] Goodman AM, Kato S, Bazhenova L, Patel SP, Frampton GM, Miller V, et al. Tumor Mutational Burden as an Independent Predictor of Response to Immunotherapy in Diverse Cancers. Molecular Cancer Therapeutics. 2017;16(11).

[64] André F, Ciruelos E, Rubovszky G, Campone M, Loibl S, Rugo HS, et al. Alpelisib for PIK3CA-Mutated, Hormone Receptor–Positive Advanced Breast Cancer. New England Journal of Medicine. 2019;380(20):1929–1940.

[65] Prasad V, Kaestner V, Mailankody S. Cancer Drugs Approved Based on Biomarkers and Not Tumor Type—FDA Approval of Pembrolizumab for Mismatch Repair-Deficient Solid Cancers. JAMA Oncology. 2017;10065.

[66] Søkilde R, Persson H, Ehinger A, Pirona AC, Fernö M, Hegardt C, et al. Refinement of breast cancer molecular classification by miRNA expression profiles. BMC Genomics. 2019;20(1):1–12.

[67] Vallon-Christersson J, Häkkinen J, Hegardt C, Saal LH, Larsson C, Ehinger A, et al. Cross comparison and prognostic assessment of breast cancer multigene signatures in a large population-based contemporary clinical series. Scientific Reports. 2019;9(1).

[68] Lundgren C, Bendahl PO, Borg A, Ehinger A, Hegardt C, Larsson C, et al. Agreement between molecular subtyping and surrogate subtype classification: a contemporary population-based study of ER-positive/HER2-negative primary breast cancer. Breast Cancer Research and Treatment. 2019;178(2):459–467.

[69] Dihge L, Vallon-Christersson J, Hegardt C, Saal LH, Häkkinen J, Larsson C, et al. Prediction of lymph node metastasis in breast cancer by gene expression and clinicopathological models: Development and validation within a population based cohort. Clinical Cancer Research. 2019;25(21):6368–6381.

[70] Persson H, Søkilde R, Häkkinen J, Pirona AC, Vallon-Christersson J, Kvist A, et al. Frequent miRNA-convergent fusion gene events in breast cancer. Nature Communications. 2017;8(1):788.

[71] Hofmann AL, Behr J, Singer J, Kuipers J, Beisel C, Schraml P, et al. Detailed simulation of cancer exome sequencing data reveals differences and common limitations of variant callers. BMC Bioinformatics. 2017;18(1):1–15.

[72] Ellrott K, Bailey MH, Saksena G, Covington KR, Kandoth C, Stewart C, et al. Scalable Open Science Approach for Mutation Calling of Tumor Exomes Using Multiple Genomic Pipelines. Cell Systems. 2018;6(3):271–281.

[73] Shi W, Ng CKY, Lim RS, Jiang T, Kumar S, Li X, et al. Reliability of Whole-Exome Sequencing for Assessing Intratumor Genetic Heterogeneity. Cell Reports. 2018;25(6):1446–1457.

[74] Costello M, Pugh TJ, Fennell TJ, Stewart C, Lichtenstein L, Meldrim JC, et al. Discovery and characterization of artifactual mutations in deep coverage targeted capture sequencing data due to oxidative DNA damage during sample preparation. Nucleic Acids Research. 2013;41(6):e67.

[75] Van Gurp TP, McIntyre LM, Verhoeven KJF. Consistent errors in first strand cDNA due to random hexamer mispriming. PLoS ONE. 2013;8(12):2–5.

[76] Danecek P, Nellåker C, McIntyre RE, Buendia-Buendia JE, Bumpstead S, Ponting CP, et al. High levels of RNA-editing site conservation amongst 15 laboratory mouse strains. Genome Biology. 2012;13(4).

[77] Piskol R, Ramaswami G, Li JB. Reliable identification of genomic variants from RNA-seq data. American Journal of Human Genetics. 2013;93(4):641–651.

[78] Horvath A, Pakala SB, Mudvari P, Reddy SDN, Ohshiro K, Casimiro S, et al. Novel insights into breast cancer genetic variance through RNA sequencing. Scientific Reports. 2013;3.

[79] Radenbaugh AJ, Ma S, Ewing A, Stuart JM, Collisson EA, Zhu J, et al. RADIA: RNA and DNA integrated analysis for somatic mutation detection. PLoS ONE. 2014;9(11).

[80] Wilkerson MD, Cabanski CR, Sun W, Hoadley KA, Walter V, Mose LE, et al. Integrated RNA and DNA sequencing improves mutation detection in low purity tumors. Nucleic Acids Research. 2014;42(13):e107.

[81] Guo Y, Zhao S, Sheng Q, Samuels DC, Shyr Y. The discrepancy among single nucleotide variants detected by DNA and RNA high throughput sequencing data. BMC Genomics. 2017;18(S6):690.

[82] Siegel MB, He X, Hoadley KA, Hoyle A, Pearce JB, Garrett AL, et al. Integrated RNA and DNA sequencing reveals early drivers of metastatic breast cancer. Journal of Clinical Investigation. 2018;128:1–13.

[83] Ramaswami G, Zhang R, Piskol R, Keegan LP, Deng P, O’Connell MA, et al. Identifying RNA editing sites using RNA sequencing data alone. Nature Methods. 2013;10(2):128–132.

[84] Banh RS, Iorio C, Marcotte R, Xu Y, Cojocari D, Rahman AA, et al. PTP1B controls non-mitochondrial oxygen consumption by regulating RNF213 to promote tumour survival during hypoxia. Nature Cell Biology. 2016;18(7):803–813.

[85] Bader AG, Kang S, Vogt PK. Cancer-specific mutations in PIK3CA are oncogenic in vivo. Proceedings of the National Academy of Sciences. 2006;103(5):1475–1479.

[86] Juric D, Rodon J, Tabernero J, Janku F, Burris HA, Schellens JHM, et al. Phosphatidylinositol 3-Kinase *α*–Selective Inhibition With Alpelisib (BYL719) in PIK3CA -Altered Solid Tumors: Results From the First-in-Human Study. Journal of Clinical Oncology. 2018;36(13):1291–1299.

[87] Avivar-Valderas A, McEwen R, Taheri-Ghahfarokhi A, Carnevalli LS, Hardaker EL, Maresca M, et al. Functional significance of co-occurring mutations in PIK3CA and MAP3K1 in breast cancer. Oncotarget. 2018;9(30):21444–21458.

[88] Yi KH, Lauring J. Recurrent AKT mutations in human cancers: consequences and effects on drug sensitivity. Oncotarget. 2015;7(4):4241–4251.

[89] She QB, Gruvberger-Saal SK, Maurer M, Chen Y, Jumppanen M, Su T, et al. Integrated molecular pathway analysis informs a synergistic combination therapy targeting PTEN/PI3K and EGFR pathways for basal-like breast cancer. BMC Cancer. 2016;16(1).

[90] Holstege H, Horlings HM, Velds A, Langerød A, Børresen-Dale AL, van de Vijver MJ, et al. BRCA1-mutated and basal-like breast cancers have similar aCGH profiles and a high incidence of protein truncating TP53 mutations. BMC Cancer. 2010;10(654).

[91] Severson TM, Peeters J, Majewski I, Michaut M, Bosma A, Schouten PC, et al. BRCA1-like signature in triple negative breast cancer: Molecular and clinical characterization reveals subgroups with therapeutic potential. Molecular Oncology. 2015;9(8):1528–1538.

[92] Pradhan MR, Siau JW, Kannan S, Nguyen MN, Ouaray Z, Kwoh CK, et al. Simulations of mutant p53 DNA binding domains reveal a novel druggable pocket. Nucleic Acids Research. 2019;47(4):1637–1652.

[93] Hisamatsu Y, Tokunaga E, Yamashita N, Akiyoshi S, Okada S, Nakashima Y, et al. Impact of FOXA1 expression on the prognosis of patients with hormone receptor-positive breast cancer. Annals of Surgical Oncology. 2012;19(4):1145–1152.

[94] Takaku M, Grimm SA, Roberts JD, Chrysovergis K, Bennett BD, Myers P, et al. GATA3 zinc finger 2 mutations reprogram the breast cancer transcriptional network. Nature Communications. 2018;9(1):1–14.

[95] Pahuja KB, Nguyen TT, Jaiswal BS, Prabhash K, Thaker TM, Senger K, et al. Actionable Activating Oncogenic ERRB2/HER2 Transmembrane and Juxtamembrane Domain Mutations. Cancer Cell. 2018;34(5):792–806.

[96] Ben-Baruch E. HER2-Mutated Breast Cancer Responds to Treatment With Single-Agent Neratinib, a Second-Generation HER2/EGFR Tyrosine Kinase Inhibitor. Journal of the National Comprehensive Cancer Network. 2015;13(9):1061–1064.

[97] Ma CX, Bose R, Gao F, Freedman RA, Telli ML, Kimmick G, et al. Neratinib efficacy and circulating tumor DNA detection of HER2 mutations in HER2 nonamplified metastatic breast cancer. Clinical Cancer Research. 2017;23(19):5687–5695.

[98] Nayar U, Cohen O, Kapstad C, Cuoco MS, Waks AG, Wander SA, et al. Acquired HER2 mutations in ER+ metastatic breast cancer confer resistance to estrogen receptor–directed therapies. Nature Genetics. 2018;51(2):207–216.

[99] Murray EM, Cherian MA, Ma CX, Bose R. HER2 Activating Mutations in Estrogen Receptor Positive Breast Cancer. Current Breast Cancer Reports. 2018;10(2):41–47.

[100] Vitting-Seerup K, Sandelin A. The Landscape of Isoform Switches in Human Cancers. Molecular Cancer Research. 2017;15(9):1206–1220.

[101] Mendoza-Villanueva D, Deng W, Lopez-Camacho C, Shore P. The Runx transcriptional co-activator, CBFbeta, is essential for invasion of breast cancer cells. Molecular Cancer. 2010;9(171).

[102] Habib JG, O’Shaughnessy JA. The hedgehog pathway in triple-negative breast cancer. Cancer Medicine. 2016;5(10):2989–3006.

[103] Locatelli M, Curigliano G. Notch inhibitors and their role in the treatment of triple negative breast cancer: Promises and failures. Current Opinion in Oncology. 2017;29(6):411–427.

[104] Weidle UH, Maisel D, Eick D. Synthetic lethality-based targets for discovery of new cancer therapeutics. Cancer Genomics and Proteomics. 2011;8(4):159–171.

[105] Hellmann MD, Ciuleanu TE, Pluzanski A, Lee JS, Otterson GA, Audigier-Valette C, et al. Nivolumab plus Ipilimumab in Lung Cancer with a High Tumor Mutational Burden. New England Journal of Medicine. 2018;378(22):2093–2104.

[106] Lauss M, Donia M, Harbst K, Andersen R, Mitra S, Rosengren F, et al. Mutational and putative neoantigen load predict clinical benefit of adoptive T cell therapy in melanoma. Nature Communications. 2017;8(1):1738.

[107] Zacharakis N, Chinnasamy H, Black M, Xu H, Lu Yc, Zheng Z, et al. Immune recognition of somatic mutations leading to complete durable regression in metastatic breast cancer. Nature Medicine. 2018;24(6):724–730.

[108] Thomas A, Routh ED, Pullikuth A, Jin G, Su J, Chou JW, et al. Tumor mutational burden is a determinant of immune-mediated survival in breast cancer. OncoImmunology. 2018;7(10):e1490854.

[109] Schmid P, Adams S, Rugo HS, Schneeweiss A, Barrios CH, Iwata H, et al. Atezolizumab and Nab-Paclitaxel in Advanced Triple-Negative Breast Cancer. New England Journal of Medicine. 2018;379(22):2108–2121.

[110] Panda A, Betigeri A, Subramanian K, Ross JS, Pavlick DC, Ali S, et al. Identifying a Clinically Applicable Mutational Burden Threshold as a Potential Biomarker of Response to Immune Checkpoint Therapy in Solid Tumors. JCO Precision Oncology. 2017;(1):1–13.

[111] Xu J, Guo X, Jing M, Sun T. Prediction of tumor mutation burden in breast cancer based on the expression of ER, PR, HER-2, and Ki-67. OncoTargets and Therapy. 2018;11:2269–2275.

[112] Shah SP, Roth A, Goya R, Oloumi A, Ha G, Zhao Y, et al. The clonal and mutational evolution spectrum of primary triple-negative breast cancers. Nature. 2012;486(7403):395–399.

[113] Cerami E, Gao J, Dogrusoz U, Gross BE, Sumer SO, Aksoy BA, et al. The cBio Cancer Genomics Portal: An open platform for exploring multidimensional cancer genomics data. Cancer Discovery. 2012;2(5):401–404.

